# decemedip: hierarchical Bayesian modeling for cell type deconvolution of immunoprecipitation-based DNA methylomes

**DOI:** 10.1101/2025.05.09.653152

**Authors:** Ning Shen, Ze Zhang, Sylvan Baca, Keegan Korthauer

## Abstract

MeDIP-seq is an enrichment-based DNA methylation profiling technique that measures the abundance of methylated DNA. While this technique offers efficiency advantages over direct methylation profiling, it does not provide absolute quantification of DNA methylation necessary for cell type deconvolution. We introduce decemedip, a Bayesian hierarchical model for cell type deconvolution of methylated sequencing data that leverages reference atlases derived from direct methylation profiling. We demonstrate its accuracy and robustness through simulation studies and validation on cross-platform measurements, and highlight its utility in identifying tissue-specific and cancer-associated methylation signatures using MeDIP-seq profiling of patient-derived xenografts and cell-free DNA. decemedip is available at https://github.com/nshen7/decemedip.

## Background

Epigenetic modifications play a pivotal role in regulating gene expression and maintaining cellular identity [1, 2]. Among the most widely studied forms of epigenetic regulation, DNA methylation (DNAme) is the addition of methyl groups primarily to cytosine bases in CpG dinucleotides (CpGs) through a dynamic process that is influenced by cell differentiation, environmental factors, aging, and disease. DNAme represents a particularly important molecular signature of cellular heterogeneity in both health and disease [3–5]. DNA methylomes obtained from mixtures of cell types such as bulk tissue samples or circulating cell-free DNA (cfDNA) generally reflect an aggregate signal that obscures the underlying cellular composition. Although experimental techniques such as cell sorting prior to DNAme profiling can provide detailed cellular information, these methods are often costly and labor-intensive. On the other hand, cell-type-specific DNAme signatures offer valuable information about distinct characteristics of individual cell types. Thus, computationally decomposing DNAme data to infer cell type proportions using such signatures [6, 7] might be essential for various applications, including cancer diagnosis, monitoring immune response, and studying developmental processes, etc [8–10].

Cell-free or bulk DNAme profiling using techniques such as methylated DNA immunoprecipitation sequencing (MeDIP-seq) [11] have emerged as a promising tool for identifying epigenetic alterations and have facilitated the discovery of noninvasive biomarkers [12–14]. MeDIP-seq is a cost-effective, affinity enrichment-based approach that selectively captures and sequences only methylated DNA fragments, providing an efficient genome-wide view of the methylome (Fig. 1A). The signals provided by MeDIP-seq are semi-quantitative measures of DNA methylation that reflect the relative abundance of methylation at the regional level (∼150–300 bp, depending on fragmentation [15]), where in general high read counts indicate regions with high methylation level and/or dense CpG context. This is in contrast to direct methylation profiling techniques such as bisulfite or enzymatic conversion, which provide absolute methylation levels (i.e. fractional levels between 0 and 1) but require the more costly whole genome sequencing, where 70–80% of the sequencing reads provide little or no information about CpG methylation [16].

**Fig. 1:**
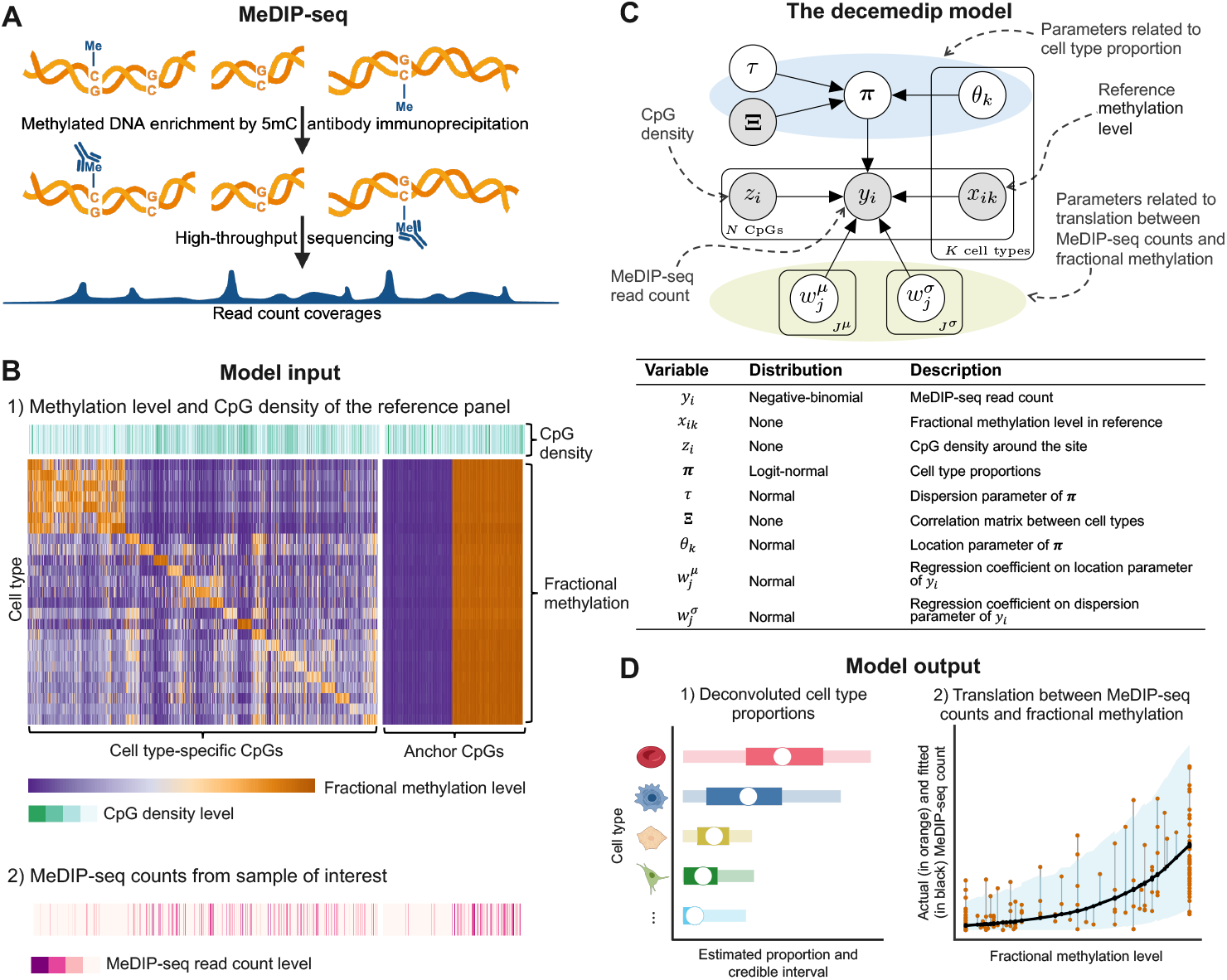
Graphical outline for decemedip. **A** Schematic representation of the MeDIP-seq experimental procedure. Methylated DNA is enriched through 5-methylcytosine (5mC)-specific antibody immunoprecipitation. The enriched DNA fragments are subsequently subjected to high-throughput sequencing, generating read count coverages that correspond to methylated regions of the genome. **B** Model input to decemedip. Part 1: Reference panel data includes cell-type-specific CpGs and anchor CpGs characterized by their CpG density (green rug) and fractional methylation levels (purpleorange heatmap). Part 2: MeDIP-seq read counts from the sample of interest (pink rug) represent the observed methylation signal. **C** Plate notation of the decemedip probabilistic graphical model. The random variables and data that form the model along with the distributional assumptions are shown. Input/observed values are denoted by gray circles. Model parameters are denoted by white circles. Please see the “Methods“ for a detailed model description. **D** Visualization of model output for the MeDIP-seq analysis. Part 1: A simple illustration of deconvoluted cell type proportions (white circles) and their credible intervals (darker colored bars represent 50% intervals and lighter colored) are displayed. Part 2: an example relationship between MeDIP-seq read coverages and fractional methylation levels modeled by decemedip. Observed MeDIP-seq counts (orange dots) are compared with the fitted curve (black line) and its 95% credible intervals (shaded blue).

While MeDIP-seq offers scalability and sensitivity, its enrichment-based nature introduces challenges for quantitative interpretation, particularly when applied to heterogeneous samples. Many statistical modeling approaches for cell type deconvolution have been developed for direct methylation profiling approaches like bisulfite sequencing and array data [17–27]. Most of these are reference-based approaches that take advantage of known methylation signatures of individual cell types from comprehensive reference atlases (i.e., large-scale, detailed collections of DNA methylation profiles across various cell types or tissues) [19, 28]. However, adapting these methodologies for data generated by enrichment-based technologies such as MeDIP-seq, requires addressing unique challenges. In particular, the most significant extrinsic challenge in developing such methodologies is the lack of similar cell-type-specific reference atlas data generated by MeDIP-seq. Existing reference atlases are predominantly derived from technologies such as whole-genome bisulfite sequencing (WGBS) or array-based platforms such as the Infinium HumanMethylation450K or EPIC BeadChip [17, 19, 21, 22, 28]. While these technologies provide comprehensive methylation profiles, they differ fundamentally from the MeDIP-seq in terms of resolution, coverage, and signal characteristics.

An intrinsic limitation of MeDIP-seq arises from its selective characterization of methylated regions. As a result, we expect that cell-type-specific hypermethylation signatures (i.e. regions that are hypermethylated in a particular cell type compared to others) would be much more informative for deconvolution than cell-type-specific hypomethylation. Nonetheless, the scarcity of hypermethylation signatures in publicly available DNAme atlases [19, 28] limits the selection of reference sites for deconvolution. In addition, as MeDIP-seq does not quantify absolute DNA methylation levels (i.e., fractional methylation), it is less effective at detecting methylation in regions with low but non-zero methylation levels and is sensitive to CpG density [29, 30]. Therefore, it is challenging to find a universal function that maps measurements between MeDIP-seq and other technologies that were used by the existing reference atlases. Previously published methods [29–31] have attempted to infer absolute methylation levels from enrichment-based read coverages while correcting for bias associated with CpG density. However, incorporating these estimated absolute levels into deconvolution analyses introduces an added layer of complexity, as it requires further calibration and handling of uncertainties arising from the estimation process, reducing the overall robustness of the resulting estimates. In our benchmarking experiments, these methods performed unsatisfactorily (results not shown), which led us to exclude them from our approach. In contrast, our model offers greater flexibility, making it better suited for the task.

These challenges require innovative computational strategies to bridge these technological gaps. In this work, we present *decemedip*, a flexible Bayesian hierarchical modeling approach specifically designed for cell type deconvolution of MeDIP-seq data, applicable to both cell-free and bulk measurements (Fig. 1). Our method represents the first DNAme-based deconvolution approach capable of incorporating cross-platform reference atlases. By integrating informative features such as CpG density, decemedip delivers an accurate and interpretable decomposition of heterogeneous DNA methylation signals.

We evaluated the performance of the proposed approach using both synthetic datasets and previously published experimental data, demonstrating its broad applicability in diverse biomedical research scenarios. Importantly, leveraging methylation profiles of cell lines with matched cross-platform measurements, we show that model components of decemedip effectively capture the relationship between MeDIP-seq counts and fractional methylation levels. Analyses of cell type deconvolution in three recent studies involving MeDIP-seq reveal that decemedip is capable of detecting unusually high proportions of cell types associated with respective cancer types or tissue samples, underscoring its utility to uncover biologically relevant insights.

## Results

### decemedip deconvolves cell types with a tailored reference panel

decemedip is a Bayesian hierarchical model designed to simultaneously infer cell type composition and characterize the relationship between fractional methylation values and MeDIP-seq read coverages. The model requires three primary inputs: a matrix of reference methylation levels derived from a comprehensive set of cell types at a curated set of CpG sites (referred to as *the reference panel*), the CpG density of genomic areas surrounding sites in the panel, and MeDIP-seq read coverages for the reference sites from the sample of interest (Fig. 1B).

Specifically, decemedip couples a logit-normal model with a generalized additive model (GAM) structure within a Bayesian framework, where the logit-normal component models cell type proportions and the GAM component models the relationship between enrichment-based read coverage and absolute DNAme levels. Our model is applied to each sample individually since the GAM model parameters may vary from sample to sample depending on variables such as sequencing depth and the form of the sample (cell-free or bulk), etc. For each site *i* ∈ {1, …, *N* } in the reference panel, the input to the model is the fractional methylation levels *x*_*ik*_ for each cell type *k* ∈ {1, …, *K*}, the CpG density level *z*_*i*_ and the MeDIP-seq read count *y*_*i*_. The logit-normal component of the model includes a unit simplex variable that follows a logit-normal prior, ***π*** = (*π*_1_, …, *π*_*K*_) where *π*_*k*_ > 0 and 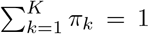. This prior describes the proportions of reference panel cell types present in the sample while taking into account the correlations between these cell types (Fig. 1B-C; Additional File 1: Fig. S1; see “Methods“ for a detailed formulation).

On the other hand, the GAM component primarily depicts the relationship between enrichment-based read coverages and fractional methylation levels derived from direct methylation profiling techniques. Specifically, a fully distributional negative-binomial model (i.e., both location and dispersion parameter are modeled with regression model) is applied to the outcome variable MeDIP-seq count *y*_*i*_, where explanatory variables include the aggregate reference methylation level across cell types, denoted as 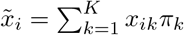, and the CpG density *z*_*i*_. Interaction terms between 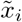 and smoothing bases of *z*_*i*_ are also included to account for the complex relationships inherent between fractional DNAme level from direct methylation profiling techniques and MeDIP-seq counts (“Methods“). We reasonably hypothesize that the regression parameters in the GAM model might differ across samples and datasets due to variation in coverage levels and batch effects, and evidence supporting the hypothesis has been presented in the case studies conducted in the article (Additional File 1: Fig. S2). For this reason, these parameters are estimated per sample.

For all experiments in this article, the selected panel of reference CpGs is specifically designed for cell type deconvolution using references derived from array-based platforms. In particular, we use MethAtlas [19], a publicly available DNA methylation atlas that compiles DNAme profiles for 25 common healthy cell types in the human body measured by 450K and EPIC arrays (“Methods“). To build a reference panel that is tailored to the characteristics of MeDIP-seq data, we first selected a curated set of CpG sites that distinguish between cell types (referred to as *cell-type-specific CpGs*; Fig. 1B).

Specifically, for each cell type, we identified CpGs that were uniquely hypermethylated relative to all other cell types. To reduce the risk of confounding cell-type-specific signals with epigenetic changes driven by cancer, potential cancer biomarkers reported by Ibrahim et al. [32] were excluded from the atlas beforehand. For each reference cell type, we selected CpGs with DNAme level that was significantly higher than the average background methylation level across all other cell types. Among these uniquely hypermethylated CpGs, the top-ranked sites based on the magnitude of difference were selected for inclusion in the reference panel (“Methods“; Additional File 2: Table S1). This approach ensures that the selected CpGs are not only cell-type-specific but also compatible with the hypermethylation-focused nature of MeDIP-seq, facilitating an accurate inference of cell type compositions.

Furthermore, unlike previously published deconvolution methods that rely solely on cell-type-specific CpGs [17–25], we additionally incorporated *anchor CpGs* into the reference panel. These CpGs exhibit relatively uniform and extreme methylation levels across cell types, enabling the algorithm to effectively and robustly capture the relationship between MeDIP-seq counts and fractional methylation values (Fig. 1B). Specifically, we selected CpG sites that are consistently highly methylated (beta values > 0.9 for all cell types in the atlas) or lowly methylated (beta values < 0.1 for all cell types) as anchor sites (“Methods“; Additional File 2: Table S2). The inclusion of anchor sites improves the identifiability of parameters and helps model the relationship at extremes, leading to a more robust posterior estimation.

Using a Bayesian formulation, decemedip infers the sample-specific posterior distributions for all model parameters, including cell type proportions (Fig. 1D, part 1). In addition, translation between MeDIP-seq counts and fractional methylation levels can be inferred based on the posteriors (Fig. 1D, part 2). The inference of posterior distributions is enabled by Markov Chain Monte Carlo (MCMC) [33–35] implementation using the Stan probabilistic programming language [36] (“Methods“).

### decemedip translates cross-platform DNAme measurements in paired cell line data

As previously described, decemedip employs a GAM component to effectively capture the relationship between fractional methylation and MeDIP-seq read counts. To evaluate the validity of this modeling choice, we used paired WGBS and MeDIP-seq data from two distinct cell lines, K562 and GM12878, acquired from the ENCODE project consortium [37–39] (Additional File 2: Table S3). Since cell lines exhibit stable characteristics across experimental conditions, they are suitable for verifying some aspects of our modeling assumptions regarding the connection between MeDIP-seq counts and fractional methylation values.

Specifically, we extracted the GAM component of decemedip and fit its Bayesian formulation to the matched cell line data using the CpGs in the reference panel (“Methods“). Fig. 2 presents the observed MeDIP-seq read coverage alongside the fitted counts predicted by the GAM component across varying fractional methylation levels and CpG density levels for the two cell lines. In both cell lines, the predicted MeDIP-seq counts closely align with the observed values, demonstrating the robustness of the GAM component in modeling the relationship between methylation levels and read counts. The 95% Bayesian credible intervals provide accurate coverage of the observed values, with coverage rates of 96.6% and 97.2% for K562 and GM12878, respectively. This indicates that the model’s uncertainty quantification was well-calibrated and effectively captured the variability in the data. The consistency of the results across varying CpG density levels underscores the model’s robustness in accommodating a variety of genomic contexts.

**Fig. 2:**
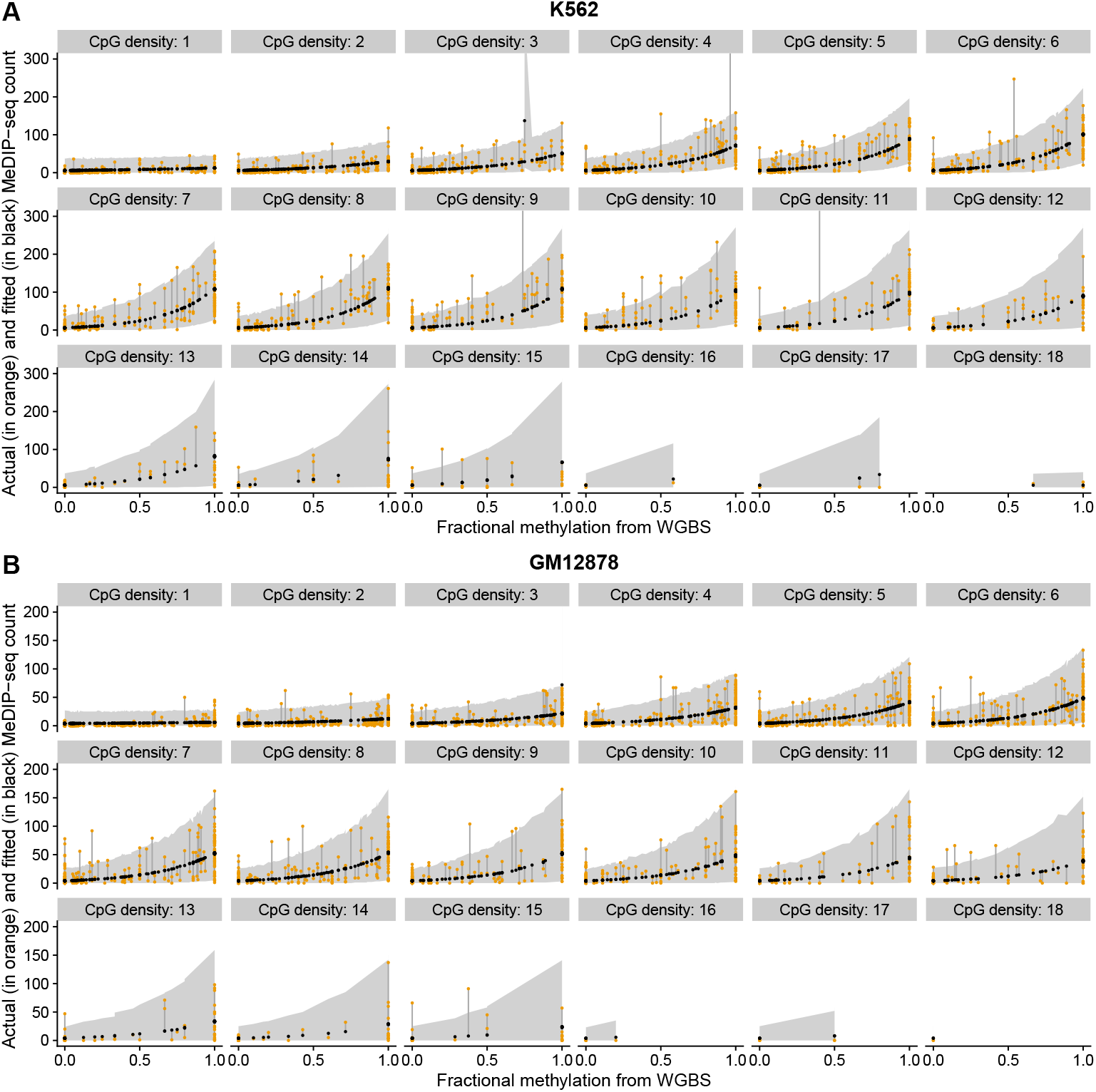
Relationship between WGBS fractional methylation and MeDIP-seq read coverage modeled by the GAM component in the decemedip model, on paired bulk cell line data. The actual MeDIP-seq read counts (orange) and the fitted counts predicted by the GAM component (black) are shown for two cell lines, **A** K562 and **B** GM12878, across varying levels of CpG density. Grey area represents the 95% credible intervals of the predicted counts. ‘CpG density: x’ means that there are x CpGs in the 100-bp window surrounding the CpG.

It is worth noting that this analysis uses WGBS data instead of the array-based data referenced in our atlas, as we were unable to find publicly available matched microarray and MeDIP-seq samples. Yet, it provides evidence to support the appropriateness of our model choice as both WGBS and array-based data reflect fractional methylation levels. Furthermore, the use of high-resolution WGBS data also underscores the flexibility and adaptability of our model to accommodate different reference atlas data types. This adaptability is crucial for extending the model’s applications across a broader range of datasets and experimental designs.

Additionally, we applied decemedip on the MeDIP-seq profiles of the two cell lines. The posterior distributions of cell type proportions are visualized in Additional File 1: Fig. S3. Notably, the majority of cell types deconvolved from GM12878 are lymphocytes including B cells (53.2%) and CD8T cells (29.5%). This result aligns with the known origin of GM12878, which was derived from B lymphocytes of a healthy female donor. In contrast, as a cell line derived from a patient with chronic myelogenous leukemia in blast crisis, the deconvolved cell type composition of K562 appears more variable and less aligned with its origin cell type. This may be attributed to cancer-related epigenomic alterations and the fact that our reference panel consists of profiles on healthy human tissues.

### decemedip identifies tissue-of-origin in bulk tumor samples

To demonstrate the ability of decemedip to identify tissue-of-origin in bulk tissue samples, we applied the method to MeDIP-seq of patient-derived xenografts (PDX) from resected metastatic prostate tumors published by Berchuck et al. [40]. PDX samples are expected to exhibit minimal within-sample heterogeneity in terms of cell type composition compared to other bulk samples, since they were selected and processed to preserve the dominant tumor cell population. Direct comparisons with alternative methods were not included, as there are no existing approaches for MeDIP-seq data.

Fig. 3 demonstrates that the estimated cell type proportions for each PDX sample indicate a dominant contribution from prostate cells, with an average percentage of 74.0% (sd = 17.3%), which is consistent with the known origins of the samples. Additionally, almost all samples (28 out of 29) display 95% credible intervals for prostate cell proportions that are distant from 0, supporting the non-zero effect with high probability (Additional File 1: Fig. S4A). The presence of colon epithelial cells was also detected, but with lower average cell type proportion (20.9%, sd = 12.3%) and fewer samples (16 out of 29) exhibited credible intervals that excluded zero compared with prostate (Fig. 3; Additional File 1: Fig. S4B). This observation may reflect the anatomical proximity of the colon to the prostate, potentially resulting from cross-contamination.

**Fig. 3:**
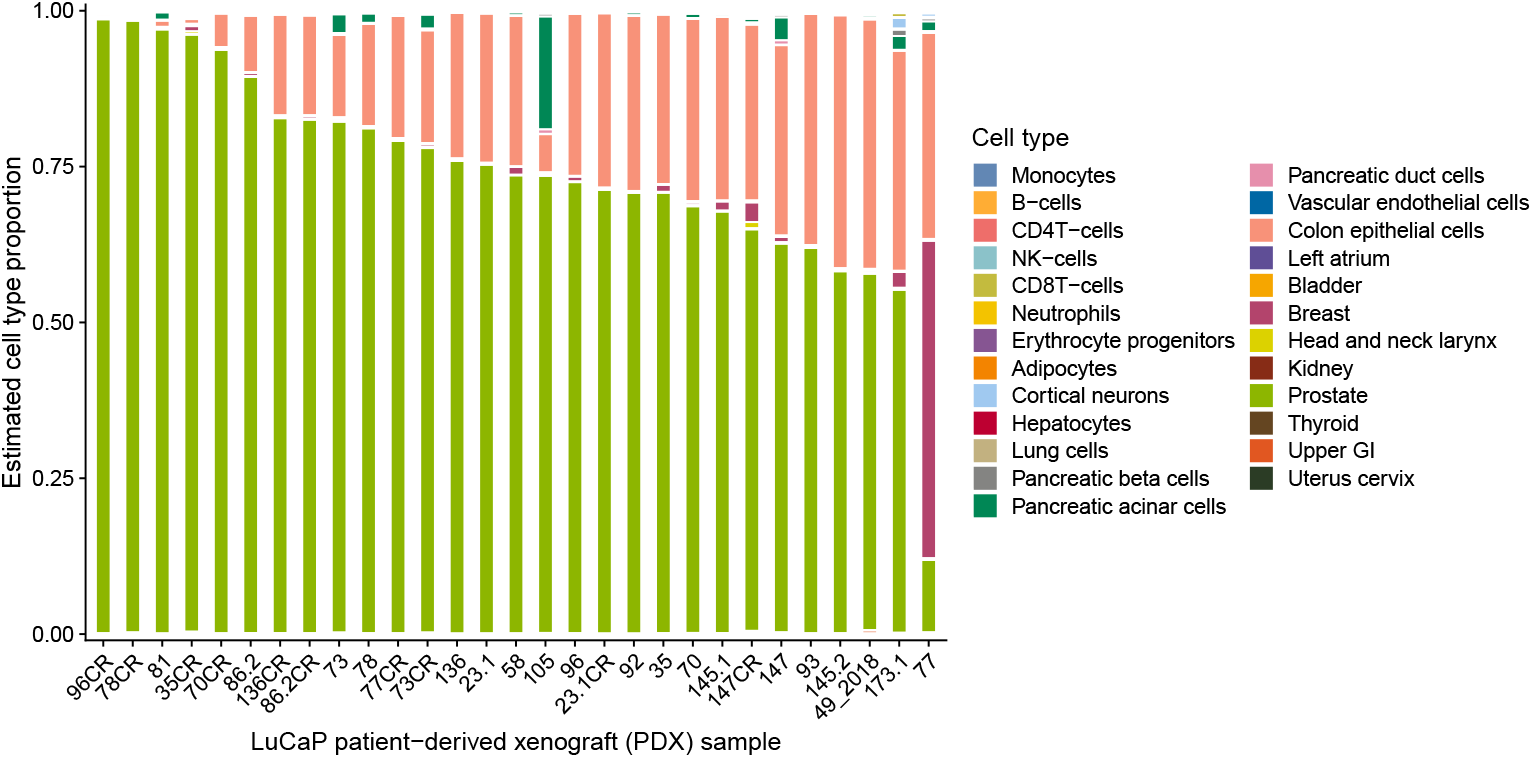
decemedip decomposition estimates for 29 patient-derived xenograft tissue samples derived from resected metastatic prostate cancer [40]. Each bar represents one xenograft tissue sample.

Although this prostate tumor cohort includes both neuroendocrine (NEPC) and adenocarcinoma (PRAD) subtypes (Additional File 2: Table S4), we did not observe pronounced neuron-associated signals in the 5 NEPC samples. On the other hand, unexpected signals from pancreatic and breast cells were detected in a small number of samples, including *105* and *77*. We speculate that these observations may stem from the absence of certain reference tissues in the atlas or artifacts caused by applying normal tissue reference to tumour cells. Such anomalies underscore the importance of careful interpretation when analyzing complex datasets.

### Benchmarking decemedip on synthetic cell-free datasets

We evaluated the performance of decemedip through in silico benchmarking experiments on synthetic sample mixtures and fully model-generated data. The two sets of experiments were designed to test the model’s ability to accurately infer model parameters from a heterogeneous mixture of cell types and assess its robustness across different data conditions.

In the first simulation study, our aim was to create mixtures of methylated cfDNA from blood cell types and varying amounts of a non-blood cell type. To do so, we computationally sampled MeDIP-seq reads originating from bulk tissues and added them to healthy cfMeDIP-seq plasma samples. Due to the limited availability of public data on healthy bulk MeDIP-seq tissue samples, we used two of the prostate tumor PDX samples from Berchuck et al. [40] that are most homogeneous in prostate cell content based on the deconvolution results (indicate range of estimated proportion prostate in these two samples). The healthy plasma samples were selected from a multicancer cell-free MeDIP-seq dataset published by Shen et al. [41] to have similar library sizes (i.e., the total number of sequenced reads, representing the depth of coverage) to the PDX samples. See “Methods“ for a detailed description of the mixture sampling procedure.

We then compared the actual percentage of prostate reads mixed in with the estimated prostate cell percentage from decemedip at mixing levels varying from 1% to 50%. Fig. 4A demonstrates that decemedip accurately inferred the prostate proportion under most settings in both sample pairs. However, some estimates showed marginal deviations from the actual proportions, with a tendency to overestimate at lower proportions and underestimate at higher proportions. This nonlinear pattern may result from discrepancies in the underlying true GAM model parameters between the PDX tissue samples and healthy plasma samples (Additional File 1: Fig. S2). However, despite these discrepancies, the posterior credible intervals were able to include the actual percentages, appropriately reflecting the uncertainty and indicating that the model provides valuable inference even under challenging conditions. As expected, hematopoietic cell types are estimated to comprise less of the mixture as we increase the ratio of read counts from PDX samples in the mixtures (Additional File 1: Fig. S5-S6).

**Fig. 4:**
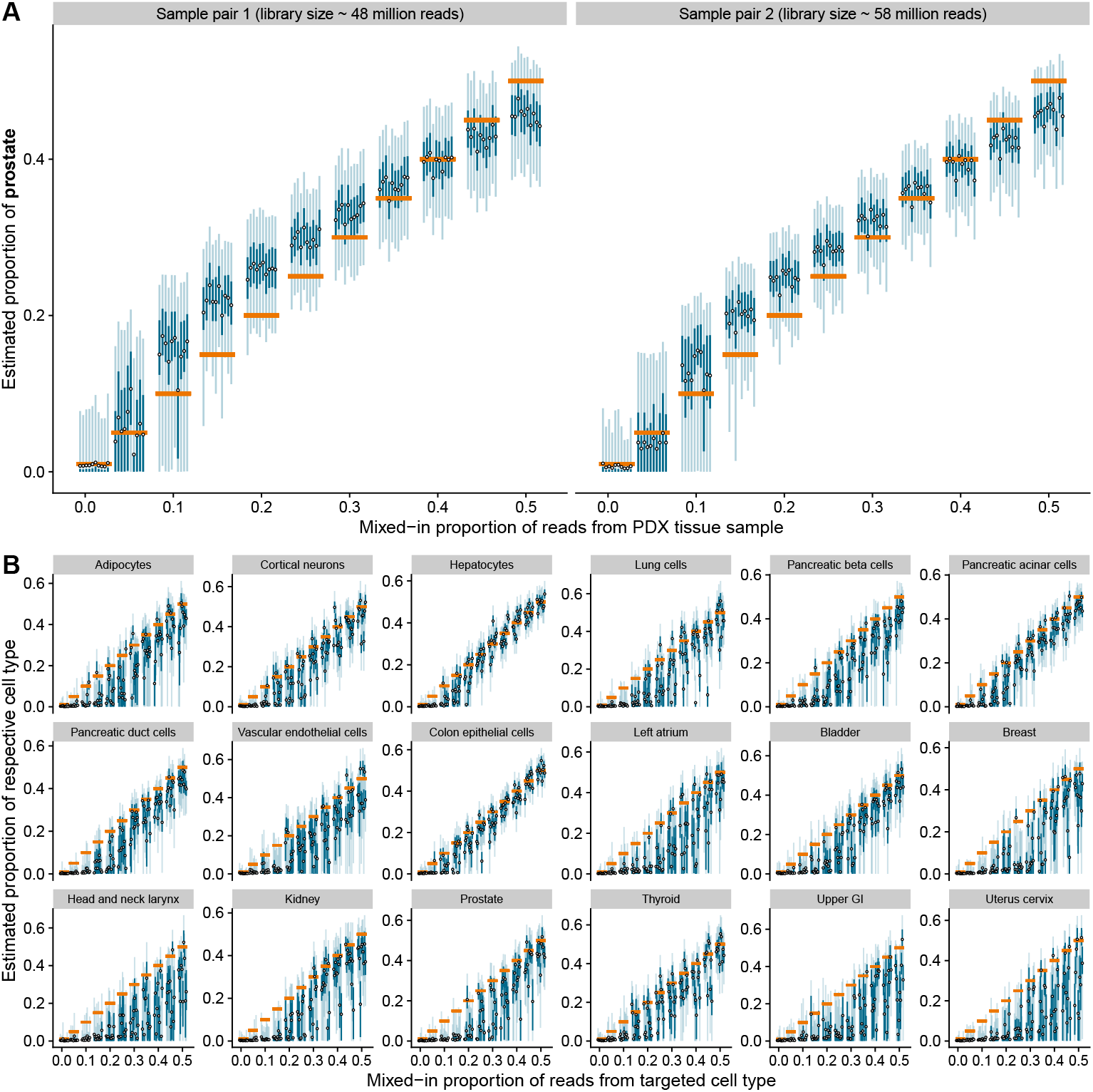
Benchmarking decemedip on synthetic cell-free MeDIP-seq read coverage. **A** Estimated prostate cell type proportions for in silico mixed samples, generated using PDX tissue samples from Berchuck et al. [40] and healthy plasma samples from Shen et al. [41]. **B** Estimated proportions for fully synthetic data generated from decemedip with varying proportions of target non-blood cell types, in a medium-coverage setting (see Additional File 1: Fig. S7 for low- and high-coverage setting). In both **A** and **B**, orange horizontal bars represent the actual proportions of the target cell types, dots indicate posterior means, and grey and blue bars represent 95% and 50% credible intervals, respectively. The percentage of targeted cell types was set from 1% to 50% per experiment. Each posterior was fitted on a synthetic sample generated by an independent random seed (“Methods“).

A second set of experiments was conducted in which synthetic read coverages were entirely generated using the decemedip model, ensuring that the ground truth of the model parameters was known. Three healthy plasma samples from Shen et al. [41] with various library sizes were selected as ‘base’ samples for determining the regression parameters. To generate the MeDIP-seq read counts, we fixed the regression parameters inferred from the base samples, set the proportions of various target cell types (from 1% to 50%), and downweighted all other cell types accordingly from their original estimated proportions. We chose to focus this experiment exclusively on nonblood cells in cell-free MeDIP-seq samples because hematopoietic cell proportions are generally less relevant in the context of cfDNA analysis and tissue-specific methylation profiles are most critical. A more detailed description of data generation and implementation is provided in the “Methods“ section.

For most target cell types, decemedip estimates of input proportions exhibited linearly increasing posterior mean estimates with increasing mixing proportions (Fig. 4B; Additional File 1: Fig. S7). However, many estimates exhibited a minor underestimation, especially breast, kidney, and prostate (Fig. 4B). This consistent bias may be attributed to inherent correlations between fractional methylation levels among the reference cell types that lead to multicollinearity during model fitting (Additional File 1: Fig. S1). The credible intervals captured the variability introduced during data generation, offering additional confidence in the estimates despite the observed bias. Such underestimation was not observed on the prostate cells in the first simulation study, which may be due to the nonlinear pattern potentially caused by discrepancies of the true parameters between the PDX tissue samples and healthy plasma samples.

Strong correlations were observed between the fitted posterior means and the actual proportions across all cell types (Additional File 1: Fig. S8). Among these, colon epithelial cells were identified with the highest agreement (corr = 0.981, *sd* = 0.010 for colon epithelial cells, 0.924 ± 0.062 for other cell types). This is likely due to a weaker association between colon epithelial cells and other cell types in the reference panel, which reduced collinearity. Furthermore, as the proportion of reads from the targeted cell type increased, the estimated regression coefficients in the GAM component remained stable and consistently fell within the 95% posterior credible intervals of the regression parameters derived from the healthy plasma samples (Additional File 1: Fig. S9-S10). These results provided evidence for the model’s appropriate prior and ability to capture the expected GAM relationship between array-based fractional methylation and MeDIP-seq read coverage.

### decemedip unveils cancer-associated cell type signatures and correlation with ctDNA proportions in cell-free data

To investigate the utility of decemedip in real-world experimental datasets, we applied the method to two multi-cancer cell-free MeDIP-seq (cfMeDIP-seq) cohorts published by Baca et al. [42] (n = 222) and Shen et al. [41] (n = 188). Both datasets include healthy individuals as controls (n = 31 and 24 in Baca et al. [42] and Shen et al. [41] respectively), serving as the baseline for interpreting the decomposition results in cancer patients. As expected, the majority of cfDNA was estimated to be hematopoietic cells in the healthy individuals for both studies (Additional File 1: Fig. S11-S13). As similarly discovered in the two publications on cell type deconvolution algorithms, Moss et al. [19] and Passemiers et al. [24], erythrocyte progenitors appeared to be one of the dominant cell types. However, the decomposition profiles from decemedip exhibited more lymphocytes and monocytes but fewer granulocytes compared to the two previously published studies. This might be attributed to the restriction of selecting cell-type-specific reference sites that are hypermethylated only for our approach, which caused stronger multicollinearity between the hematopoietic cell types in the reference panel compared to the previous studies.

Fig. 5 displays the deconvoluted proportions of cell types that are specifically associated with cancer types in the Baca et al. [42] dataset. Patients exhibited significantly elevated proportions of their respective tissue-specific cell types compared to healthy controls, especially in the patient group of breast, colorectal, and prostate cancer.

**Fig. 5:**
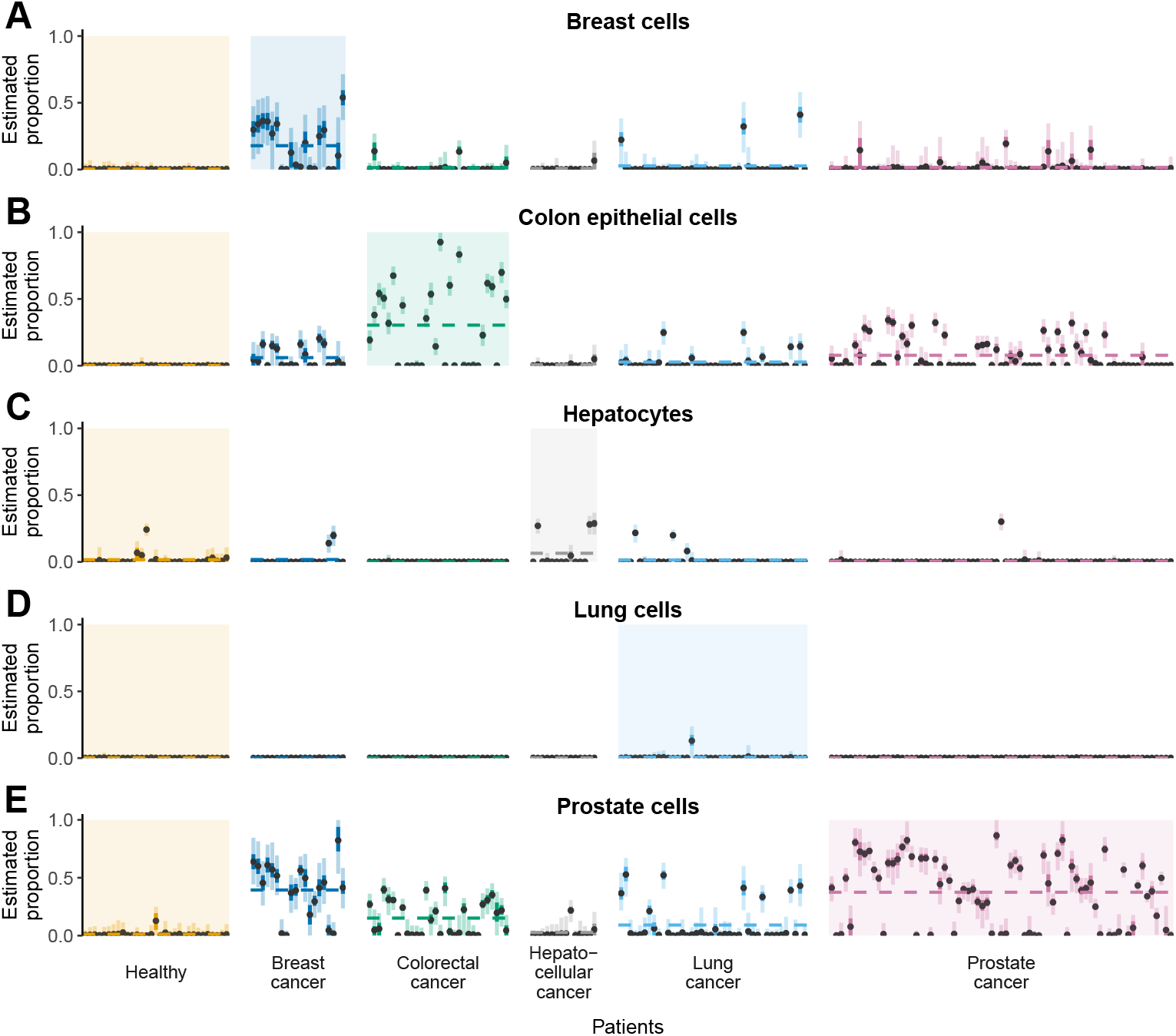
Deconvoluted cell type proportions in cancer patients in Baca et al. [42]. The posterior distributions of cell type proportions for **A** breast cells, **B** colon epithelial cells, **C** hepatocytes, **D** lung cells, and **E** prostate cells are presented for healthy controls and cancer patients. These five cancer types were selected because each is uniquely associated with a single cell type in the reference panel. Each interval represents the fitted posterior distribution for the cell type proportion of an individual patient, categorized and colored by patient groups. Within each interval, the dots denote the posterior mean, while the light and dark bars indicate the 95% and 50% credible intervals, respectively. Highlighted background colors emphasize the group comparisons of interest for the respective cell types. Horizontal dashed lines indicate the average posterior means of the cell type proportions within each patient group.

Such findings highlight the ability of decemedip to correctly detect cfDNA from cancer tumors. On the other hand, we notice that some patients also exhibited increased proportions of cell types irrelevant to their cancer types, for example, prostate cells in breast cancer patients. Similar anomalies involving cancer patients were observed in other studies that focus on cell type deconvolution [19, 24]. We speculate that these unexpected estimates might be attributed to cancer-related epigenomic alterations. We note that our reference panel is composed of healthy cell type samples. Although we have made an effort to remove tumor-related biomarkers from the reference panel (“Methods“), it is unlikely that all cancer-specific epigenetic alterations have been eliminated and residual tumor-associated methylation changes may still be present.

Analogous findings were observed in the Shen et al. [41] study, though non-zero proportions of tumor-associated cell types were estimated from fewer patient samples compared to the Baca et al. [42] dataset (Additional File 1: Fig. S14). However, significantly higher proportions of B-cells and CD8T-cells were observed in the acute myeloid leukemia (AML) patients compared with healthy controls and other cancer patients, potentially reflecting an immune response that is unique to AML (Additional File 1: Fig. S13 & S15).

Notably, in addition to identifying cancer-specific cell type signatures, decemedip revealed significant increases in non-blood cell proportions in cancer patients compared to healthy controls across most patient groups in both case studies (Fig. 6A and Additional File 1: Fig. S16). This consistent elevation suggests that non-blood cell-derived cfDNA serves as a robust indicator of cancer presence.

**Fig. 6:**
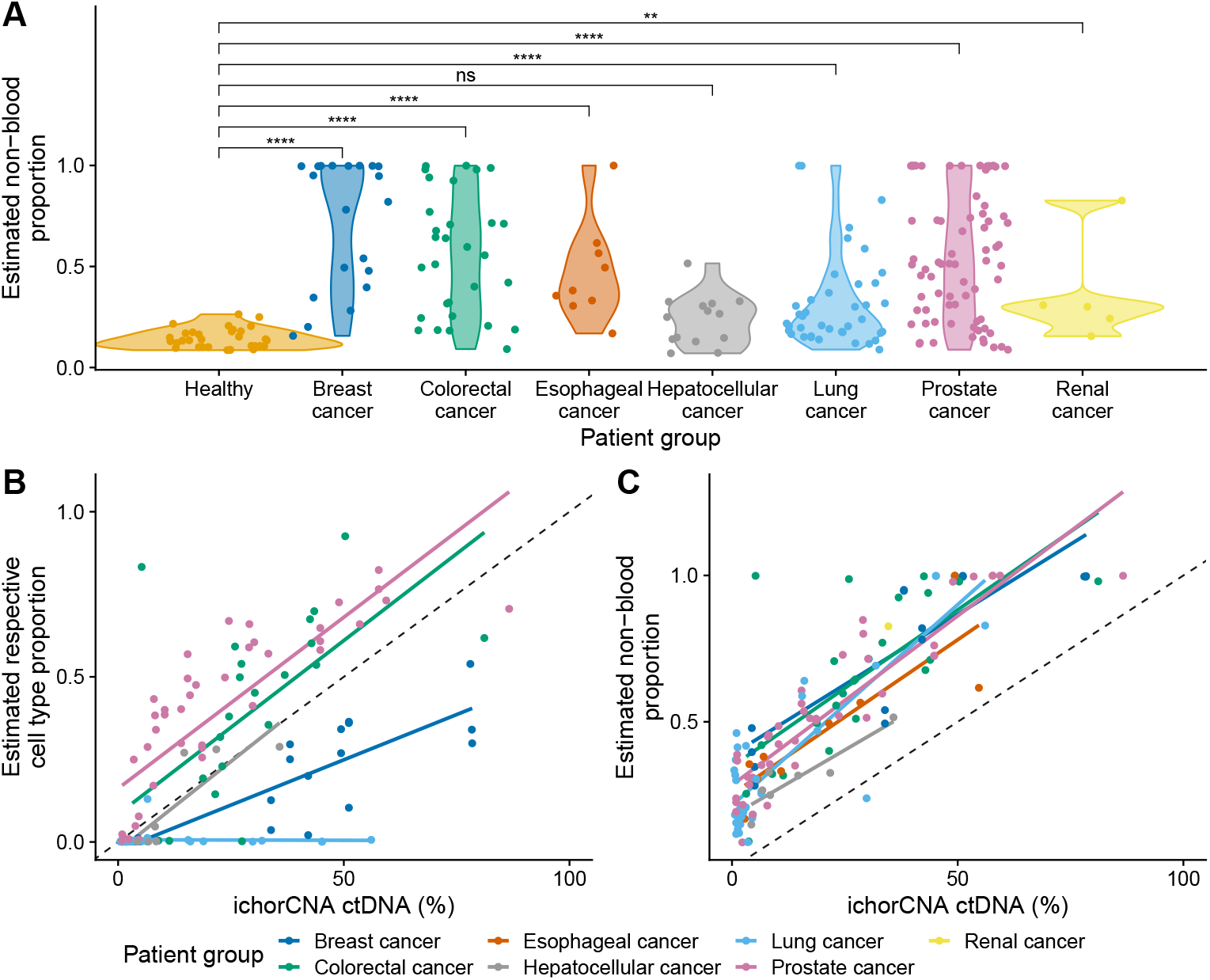
Estimated proportions of non-blood and tissue-specific cell types across patient groups and ctDNA Levels in Baca et al. [42]. **A** Estimated total non-blood cell proportions in all patient groups and comparison between cancer groups and the healthy controls. The number of stars represents the significance level of Bonferroni-corrected p-values from two-sample comparisons conducted via Wilcox tests (****:< 10^−4^, ***: < 0.001, **:< 0.01, *:< 0.05, ns: not significant). **B** Relationship between ichorCNA-estimated ctDNA levels and the estimated proportions of respective cell types corresponded to cancer types (breast cancer: breast, lung cancer: lung, colorectal cancer: colon epithelial cells, hepatocellular cancer: hepatocytes, prostate cancer: prostate). Only cancer types that exclusively associate with a single cell type in the reference panel were included. Only samples with non-zero ctDNA levels were included. **C** Relationship between the ctDNA levels and the overall estimated non-blood cell type proportion. For **A-C**, each dot represents a patient, colored by patient groups; and for **B-C**, the straight lines were fitted by linear regression with respect to each patient group.

Last but not least, the Baca et al. [42] study also provides circulating tumor DNA (ctDNA) content inferred from low-pass whole-genome sequencing data using ichorCNA [43], which serves as a valuable independent measure to validate the cell type proportion estimates of decemedip (“Methods“). Fig. 6B showcases strong positive correlations between cancer-associated cell type proportions and ctDNA levels across all cancer types. Lung cells were an exception due to their low proportion estimates. However, non-blood proportions in all cancer types, including lung, exhibited strong correlations with ctDNA levels (Fig. 6C), reinforcing the biological relevance of decemedip’s estimates and its value as a diagnostic and research tool.

## Discussion

Enrichment-based DNA methylation sequencing technologies, such as MeDIP-seq, offer scalable and cost-effective methylome profiling across diverse samples, creating a growing need for methods capable of robustly estimating cell type composition while addressing biases inherent to these technologies. To tackle the challenges of selective enrichment, sparsity, and lack of compatible reference data, we introduced decemedip, a Bayesian hierarchical framework for cell type deconvolution. Unlike earlier methods designed for bisulfite sequencing or array-based platforms, which assume direct measurements of absolute methylation levels, decemedip leverages a flexible probabilistic model that integrates CpG site-specific information from methylated sequencing and alternative reference platforms. This enables accurate estimation of cell type proportions from MeDIP-seq data while accounting for enrichment-based signals that do not reflect single-CpG resolution but rather represent the relative abundance of methylation-enriched DNA fragments.

Using paired cell line data, we demonstrated the model’s ability to capture the relationship between fractional methylation levels measured with arrays and MeDIP-seq read counts, achieving high accuracy and well-calibrated uncertainty estimates across CpG density levels (Fig. 2). This validation of our modeling choices in the GAM component highlights its robustness in handling enrichment-based signal variability associated with CpG density. Beyond controlled cell line experiments, applying decemedip to patient-derived xenograft samples provided evidence of its utility in real-world bulk samples. The model effectively identified expected tissue-specific cell types and highlighted unexpected signals, such as potential cross-contamination from neighboring tissues (Fig. 3).

To further demonstrate that decemedip is robust across diverse scenarios, we conducted simulation studies that generate realistic cell-free mixtures. We observed that decemedip was able to accurately estimate cell type proportions under varying coverage and mixture conditions in these simulations (Fig. 4; Additional File 1: Fig. S7). This suggests that the model is well-equipped to handle the inherent variability in sequencing depth and sample composition, making it broadly applicable across different experimental settings. Additionally, analyses of cfMeDIP-seq of patient samples demonstrate decemedip’s effectiveness in uncovering cancer-associated cell type signatures and detect meaningful correlations with ctDNA levels (Fig. 5-6). These findings suggest that decemedip could potentially serve as a valuable tool for non-invasive cancer detection and monitoring by identifying tumor-derived signals within cell-free DNA. Finally, it is worth noting that while our case studies focused on MeDIP-seq, the model’s flexibility suggests it could be adapted to other enrichment-based platforms, such as MBD-seq [44] and MRE-seq [45], with appropriate modifications to the reference panel.

Some aspects of our proposed approach could benefit from additional improvement. First, incorporating a more comprehensive cell type reference atlas, including a broader array of cell types, would enable more precise deconvolution, particularly for complex tissues or samples with rare cell populations. Second, we currently consider a flat correlation structure between the cell types, while integrating cell type lineage into the model could allow it to account for complex relationships between cell types and might help to distinguish closely related cell types. Finally, non-CpG methylation and cfDNA fragment length might offer complementary information about cell types of origin, thus expanding the scope of input data to include this information could enhance the model’s ability to capture distinct epigenetic and biophysical characteristics of each cell type.

## Conclusions

We introduced decemedip, a Bayesian hierarchical modeling approach with a software implementation specifically designed to perform cell type deconvolution from MeDIP-seq data, addressing challenges unique to enrichment-based DNA methylation data.

To the best of our knowledge, this is the first deconvolution method that is applicable to affinity enrichment data and capable of leveraging a reference panel derived from alternative platforms. decemedip estimates cell type proportions with accurate uncertainty quantification while overcoming the challenges inherent to MeDIP-seq, including selective enrichment and sparsity. Moreover, since MeDIP-seq is sensitive to CpG density, we designed decemedip to be robust to this variation, offering a flexible framework for diverse biological samples and contexts.

We extensively evaluated the performance of decemedip using paired cross-platform cell line data, in silico mixtures, simulations, and real-world applications involving patient-derived xenografts and cfDNA from cancer patients. In the in silico mixtures and simulation studies, the method demonstrated high accuracy in estimating cell type proportions, providing well-calibrated uncertainty estimates even under varying coverage and mixture conditions. In the case studies, decemedip successfully identified tissue-specific signatures and revealed subtle patterns of cell type composition in heterogeneous samples. We also discovered strong associations between the composition estimates and independently measured tumour ctDNA level, further validating decemedip’s biological relevance.

In summary, decemedip is a robust and innovative solution for cell type deconvolution in enrichment-based methylation data, bridging the gap between MeDIP-seq and alternative methylomic platforms. It offers a reliable framework for uncovering tissuespecific and disease-associated methylation patterns, serving as a valuable tool for advancing both basic research and the potential of translational applications in epigenomics. By leveraging the efficiency of MeDIP-seq, decemedip provides a cost-effective and scalable alternative to whole-genome bisulfite sequencing while maintaining high accuracy in deconvolution tasks. This efficiency makes it particularly well-suited for large-scale studies and clinical applications where sequencing depth and cost are limiting factors.

## Methods

### The decemedip model

#### Model inputs

The decemedip model requires a reference dataset containing fractional methylation levels (such as from microarray or WGBS), composed of *K* cell types indexed by *k*, at *N* CpG sites indexed by *i*. The fractional methylation levels from specific cell types and a reference CpG set are collected in a *N* × *K* matrix: **X** = (*x*_*ik*_)_1≤*i*≤*N*,1≤*k*≤*K*_, where *x*_*ik*_ is the fractional methylation level on the site *i* for cell type *k*. Additionally, the model requires the CpG density levels **z** = (*z*_*i*_)_1≤*i*≤*N*_ for the reference CpG sites. Specifically, the density level is defined as the log1p-transformed count of CpG sites within a 100-bp window that centers at the reference site. Together, **X** and **z** represent the reference panel information. For sample data, the model takes an individual sample sequenced by MeDIP-seq at a time. It shall be provided with the enrichment-based read coverages from the reference CpGs, **y** = (*y*_*i*_)_1≤*i*≤*N*_, extracted from the sample of interest.

#### Generative process of the model

As mentioned in the main text, the decemedip model primarily consists of two components: a hierarchical logit-normal model that represents the cell type proportions, and a GAM component that reflects the relationship between enrichment-based read coverages and fractional methylation levels adaptable to various CpG-density contexts. We assume that the cell type proportion of the MeDIP-seq sample ***π*** = (*π*_1_, …, *π*_*K*_) follows a logit-normal distribution:

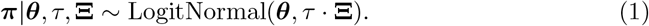

In Eq. 1, ***θ*** = (*θ*_1_, …, *θ*_*K*_) is the mean vector in the pre-softmax space representing the location parameter. The parameter **Ξ** is a correlation matrix of dimension *K* × *K* that introduces associations between the components of the logit-transformed vector (i.e., the proportions of cell types). The parameter *τ* is a scalar that scales the correlation matrix, which acts as a precision parameter, influencing the variability of ***π***. Parameters *τ* and **Ξ** together determine the shape and spread of the distribution in the logit space.

Assuming the cell type proportion vector ***π*** of a MeDIP-seq sample has been generated, we may obtain the *underlying absolute methylation levels*, 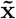, from the sample of the reference sites by aggregating from the reference absolute mathylation levels:

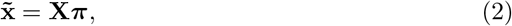

Or

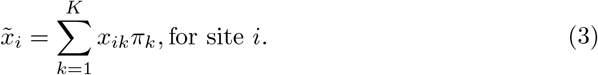

Then, we assume a distributional model on the enrichment-based read coverages under the GAM framework:

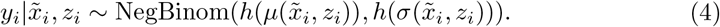

The negative-binomial distribution was used to account for the data overdispersion. In Eq. 4, *µ*(·, ·) and *σ*(·, ·) are mean and dispersion functions, which are transformed via the link function *h*(·) in the negative binomial model. We adopted the exponential function as the link function throughout the study, i.e., *h*(·) = exp(·).

The mean and dispersion are expanded as linear combinations of basis functions *b*_*j*_(*z*_*i*_) and predictors 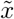, defined as:

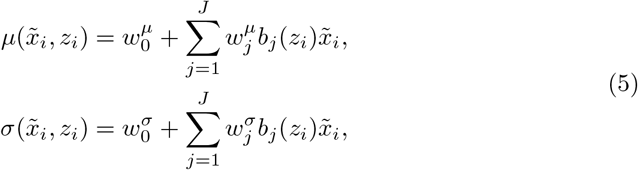

In Eq. 5, *b*_*j*_(·) are the basis functions of **z** including an intercept term, and 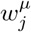 and 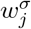 for *j* ∈ {1, …, *J*}, are the regression coefficients corresponding to the mean and dispersion functions respectively, where *J* is the total number of basis functions applied to **z**. Interactions between bases of *z* and aggregate methylation level 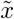 are included in the regression model to account for the complex relationship between fractional methylation and MeDIP-seq counts that is subject to the change of CpG-density. We adopted the cubic polynomial bases for *z* throughout experiments in this article, i.e. *b*_1_(*z*) = 1, *b*_2_(*z*) = *z, b*_3_(*z*) = *z*^2^, *b*_4_(*z*) = *z*^3^, *J* = 4 in this case. Though simple, the complexity of cubit polynomial bases already suffice for our model due to limited data points (Fig. 2). However, more complicated forms of the bases such as B-splines are available in our software implementation, which might be applicable if applied to an expanded reference panel.

#### The full likelihood

The full likelihood of decemedip can be formulated as:

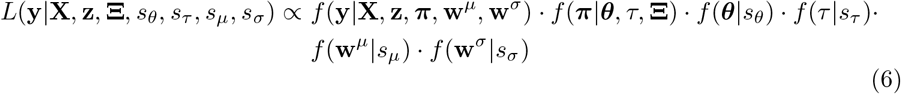

where 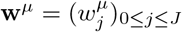 and 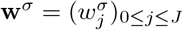. On the right-hand side of this likelihood function, the first term represents the GAM component, the second term corresponds to the logit-normal component, and the remaining terms are the prior distributions on the latent parameters.

In addition, we applied a weighted likelihood approach to reduce the influence of anchor sites relative to cell type-specific sites. This weighting strategy ensures that the model prioritizes the cell-type-specific sites, which carry the key information for estimating proportions, while still incorporating the anchor sites to capture extreme-value behavior without overwhelming or distorting the estimates. By downweighting anchor sites, we avoid potential overemphasis on their contributions, which might otherwise lead to biased or unstable estimates, given their secondary role in the modeling process. The formal formulation of the log-likelihood is as follows:

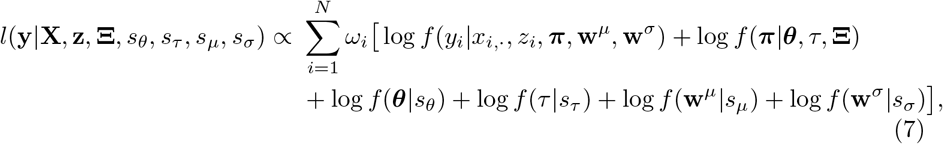

In all experiments presented in this article, we applied

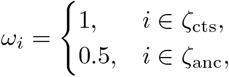

where ζ_cts_ and ζ_anc_ denote the index sets of cell-type-specific CpGs and anchor CpGs, respectively.

#### Prior choices

In the logit-normal component, each *θ*_*k*_ is assumed to be independently distributed with a prior *θ*_*k*_ ∼ 𝒩 (0, *s*_*θ*_), for *k* ∈ {1, …, *K*}. The parameter *τ* also follows a normal prior *τ* ∼ 𝒩 (0, *s*_*τ*_). The parameter **Ξ** is considered fixed and is empirically estimated by the pairwise correlation matrix between cell types using the cell-type-specific sites in the reference panel **X** in all experiments. In the GAM component, each 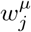 and 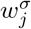 is assumed to be independently distributed with a normal prior, i.e., 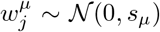 and 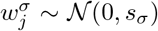, for *j* ∈ {0, …, *J* }. In all experiments presented in this article, all dispersion parameters in the prior distributions are set to 3, i.e., *s*_*θ*_ = *s*_*τ*_ = *s*_*µ*_ = *s*_*σ*_ = 3. These hyperparameters were chosen to represent a mildly informative prior, ensuring minimal regularization while maintaining the stability of MCMC sampling.

#### Implementation

The posterior distribution for the model parameters in decemedip is not amenable to analytic solutions. Hence, we resort to MCMC [33–35] implementations using the Stan probabilistic programming language [36]. In all experiments presented in this article, we utilized MCMC instead of variational Bayes methods because high accuracy is critical for our inference, and the computational cost is controllable and acceptable due to a relatively low-dimensional model and small sample sizes. decemedip is publicly available as an R package at https://github.com/nshen7/decemedip and has been submitted to Bioconductor.

#### The reference panel

##### MethAtlas

To construct our reference panel, we used the genome-wide DNA methylation atlas, MethAtlas, published by Moss et al. [19]. MethAtlas is a comprehensive database of DNA methylation consisting of 25 key human cell types, including only primary healthy tissues. Some of these cell type methylation profiles were collected from public consortiums such as The Cancer Genome Atlas (TCGA) and the Gene Expression Omnibus (GEO). The authors generated the others by isolating primary cells via flow cytometry or magnetic beads. All genome-wide methylomes were analyzed using Illumina 450K or EPIC BeadChip array platforms in MethAtlas. MethAtlas was chosen because it provides a relatively sufficient number of hypermethylated cell-type-specific sites, compared to the other publicly available DNAme atlases such as the Loyfer et al. [28].

##### cell-type-specific CpGs

To minimize the potential overlap between cell-type-specific signals and cancer-related epigenetic changes, we first excluded probes from MethAtlas that lie in differentially methylated regions (DMRs) associated with various tumor types (as listed in Supplementary Table 4 of Ibrahim et al. [32]). We denote *β*_*ik*_ as the beta values for site *i* and cell type *k* in MethAtlas. The difference of beta values between the target cell type *k* and the average of other cell types is defined as 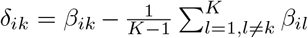. For each cell type, we selected the top 100 CpGs ranked by*δ*_*ik*_ in descending order from the subset of CpGs that exhibit maximum beta values in the target cell type *k*. Therefore, this procedure identified a total of 2,500 19 cell-type-specific CpG sites. The CpG information and beta values these reference sites can be found in Additional File 2: Table S1.

##### Anchor CpGs

Anchor CpGs were selected from CpG sites that consistently exhibit extreme beta values across all cell types. Specifically, a CpG site *i* is defined as all-tissue methylated if *β*_*ik*_ > 0.9, and as all-tissue unmethylated if *β*_*ik*_ < 0.1 for all *k* ∈ {1, …, *K*} . From each of these categories, 500 CpG sites were selected to constitute the set of anchor CpGs. We sampled the anchor CpGs so that the distribution of CpG density (i.e., number of CpGs in their surrounding 100-bp windows) resembles the distribution in cell-type-specific CpGs. Consequently, this procedure resulted in 1,000 anchor CpG sites, and hence a total of 3,500 sites in the reference panel. The CpG information and beta values these reference sites can be found in Additional File 2: Table S2.

### Paired cell line dataset

#### Data processing

WGBS and MeDIP-seq data for the two cell lines (K562 and GM12878) were obtained from the ENCODE project consortium [37–39], with data identifiers detailed in Additional File 2: Table S3. WGBS files were downloaded in BED format, and their genomic coordinates were converted from hg38 to hg19 using liftOver [46]. MeDIP-seq files were downloaded in BAM format aligned to the hg19 genome assembly.

#### The GAM model fitted to cell line data

The exact same model structure of the GAM component in the decemedip model was applied to the paired cell line data, except that the aggregate methylation level 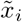 is replaced by the fractional methylation measured by WGBS,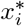:

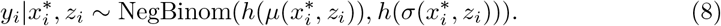

Where 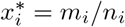, with *m*_*i*_ and *n*_*i*_ being the number of methylated reads and the total number of reads covering site *i* in the WGBS profile. The model was implemented in Stan and the posterior distribution was inferred using MCMC as well. CpG sites with no coverage from WGBS were removed during model fitting, resulting 159 (4.5%) and 94 (2.7%) out of 3,500 CpGs in the reference panel removed for K562 and GM12878, respectively.

#### Prostate cancer xenograft dataset

MeDIP-seq was performed on 29 LuCaP PDX samples, including 5 from neuroendocrine prostate cancer (NEPC) and 24 from castration-resistant prostate adenocarcinoma (PRAD). Details on quality control and data pre-processing on raw reads can be found in Berchuck et al. [40]. See Additional File 2: Table S3 for sample metadata. Fastq files were provided by the authors of Berchuck et al. [40]. Trim Galore (v0.6.10, https://github.com/FelixKrueger/TrimGalore) was used to remove adapters and trim poor-quality sequencing reads with default parameters in paired-end mode. After trimming, the reads were aligned to the human reference genome using bowtie2 (v2.4.1) [47, 48] with default parameters in paired-end mode. Human genome (hg19/ GRCh37) was downloaded from the University of California Santa Cruz (UCSC) genome browser (https://genome.ucsc.edu/). SAMtools (v1.16.1) [49] with default settings was used to convert SAM to BAM format, filter out duplicates, sort and index the files, and provide mapping statistics for the output.

### Multi-cancer cell-free datasets

#### The Baca et al. [42] dataset

In total n = 222 plasma cfMeDIP-seq samples were included in our experiments, including 31 from healthy donors, 20 from breast cancer patients, 30 from colorectal cancer patients, 9 from esophageal cancer patients, 14 from hepatocellular cancer patients, 40 from lung cancer patients, 73 from prostate cancer patients and 5 from renal cancer patients. Reads were aligned to the hg19 human genome. Details on the quality control and data processing can be found in Baca et al. [42]. Please also refer to Baca et al. [42] for details regarding ctDNA estimation.

#### The Shen et al. [41] dataset

In total n = 188 plasma cfMeDIP-seq samples were included in our experiments, including 24 from healthy donors, 27 from AML patients, 20 from bladder cancer (BL) patients, 25 from breast cancer (BRCA) patients, 23 from colorectal cancer (CRC) patients, 25 from lung cancer (LUC) patients, 24 from pancreatic cancer (PDAC) patients and 20 from renal cancer (RCC) patients. Details on the quality control and data processing can be found in Shen et al. [41]. Processed BAM files were provided by the authors upon approval of a Data Transfer Agreement.

#### Generating synthetic datasets

#### Synthetic sample mixtures

Two pairs of samples from the prostate tumor PDX dataset [40] and the cfMeDIP-seq dataset published by Shen et al. [41] were used to generate synthetic mixtures. The first pair included *78CR* (tumor tissue) and *Control 7* (healthy plasma) with a sequencing coverage of approximately 48 million reads, while the second pair consisted of *81* (tumor tissue) and *Control 14* (healthy plasma) with a coverage of approximately 58 million reads. Samples *78CR* and *81* were estimated to contain 98.1% and 96.9% of prostate cells, respectively. For each pair, ten synthetic mixtures were generated at varying tissue mixed-in proportions *ρ*, where *ρ* ∈ {0.01, 0.05, 0.1, 0.15, 0.2, …, 0.5} .

The synthetic mixtures were constructed as follows: First, the total read coverage for the reference CpGs in both the healthy plasma and PDX tissue samples was calculated and denoted as *C*^plasma^ and *C*^tissue^, respectively. Reads were then subsampled from the original experimental distributions using a binomial model. Specifically, for a given CpG site *i*, denoting the read count on the reference CpG site *i* from the plasma and tissue sample as 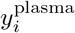 and 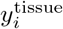. The number of reads originating from plasma and tissue is sampled from Binom(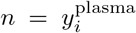, *p* = 1 − *ρ*) and Binom(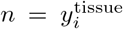, *p* = *ρ* · *C*^plasma^*/C*^tissue^), ensuring that the expected read ratio between plasma and tissue in the synthetic sample was 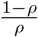. Given a *ρ*, the expected proportion of prostate is hence 0.981 · *ρ* and 0.969 · *ρ* for the two sample pairs respectively.

#### Model-generated data

The three ‘base’ healthy plasma samples from Shen et al. [41] are *Control 24, Control 3*, and *Control 14*, with library sizes of 2.1 × 10^7^, 4.4 × 10^7^, and 5.8 × 10^7^ reads, respectively. First, to match the characteristics of observed experimental samples across varying coverage levels, the decemedip model was fitted to each plasma sample to estimate regression parameters and blood cell type proportions. Using the fitted parameters from each base sample, ten synthetic samples were generated with a mixed-in proportion of a target cell type, *ρ*, where *ρ* ∈ {0.01, 0.05, 0.1, 0.15, 0.2, …, 0.5} . Specifically, for a given target cell type and mixed-in proportion *ρ*, synthetic read counts were generated by assigning the target cell type a proportion of *ρ* and proportionally downweighting all other cell types to sum to 1 − *ρ*. Default prior choices were applied to the generative process.

## Acronyms

AML: acute myeloid leukemia. 14, 21
cfDNA: cell-free DNA. 2, 9, 11, 12, 16
cfMeDIP-seq: cell-free MeDIP-seq. 11, 21
ctDNA: circulating tumor DNA. 14
DMR: differentially methylated region. 19
GAM: generalized additive model. 5, 6, 9, 11, 15, 17–20
GEO: Gene Expression Omnibus. 19
MCMC: Markov Chain Monte Carlo. 6, 19, 20
MeDIP-seq: methylated DNA immunoprecipitation sequencing. 2–7
PDX: patient-derived xenografts. 8, 10, 11, 20, 21
TCGA: The Cancer Genome Atlas. 19
WGBS: whole-genome bisulfite sequencing. 2, 6–8, 16, 20

## Declarations

## Ethics approval and consent to participate

Ethics approval was granted for this study by the UBC Clinical Research Ethics Board (H24-00478).

## Consent for publication

Not applicable.

## Availability of data and materials

WGBS and MeDIP-seq data for the two cell lines, K562 and GM12878, were obtained from the ENCODE project consortium (Additional File 2: Table S3). The prostate cancer xenograft dataset is publicly available at GSE290990. The multi-cancer cell-free datasets [41, 42] shall be available from the corresponding authors upon reasonable request. The decemedip framework is publicly available as R software released under the MIT license (see https://github.com/nshen7/decemedip [50]). A snapshot of the software used for generating results in this manuscript can be found at https://doi.org/10.5281/zenodo.15186373 [51]. The code used to process and analyze the data is available at https://github.com/nshen7/decemedip-experiments [52] under the MIT license, and a snapshot is available at https://doi.org/10.5281/zenodo.15186380 [53].

## Authors’ contributions

NS and KK conceptualized the problem. NS and KK developed the methodology. NS implemented the method and wrote the R package. NS conducted full or partial of all studies and drafted the manuscript. ZZ contributed to the deconvolution analyses of cell lines and Baca et al. [42] dataset. NS, KK, ZZ, and SB contributed to reviewing and editing the manuscript. All authors have approved the final manuscript.

## Acknowledgements

This research was supported in part through the computational resources and services provided by Advanced Research Computing at the University of British Columbia. We thank Dr. Matthew Freedman for facilitating access to the multi-cancer cohort from Baca et al. [42].

If any of the sections are not relevant to your manuscript, please include the heading and write ‘Not applicable’ for that section.

## Additional Files

**Additional File 1 (Additional File 1.pdf)**: Supplementary materials.

**Additional File 2 (Additional File 2.xlsx)**: Table S1-S2.

**Table S1:** cell-type-specific CpG sites in reference panel. Genomic coordinates were mapped to hg19.

**Table S2:** Anchor CpG sites in reference panel. Genomic coordinates were mapped to hg19.

**Table S3:** ENCODE data sources of cell lines used in the experiments.

**Table S4:** Sample metadata in prostate cancer xenograft dataset from Berchuck et al.

Editorial Policies for:

Springer journals and proceedings: https://www.springer.com/gp/editorial-policies

Nature Portfolio journals: https://www.nature.com/nature-research/editorial-policies

*Scientific Reports*: https://www.nature.com/srep/journal-policies/editorial-policies

BMC journals: https://www.biomedcentral.com/getpublished/editorial-policies

