## Additional File 1 for "decemedip: hierarchical Bayesian modeling for cell type deconvolution of immunoprecipitation-based DNA methylomes"

### S1 Supplementary Figures

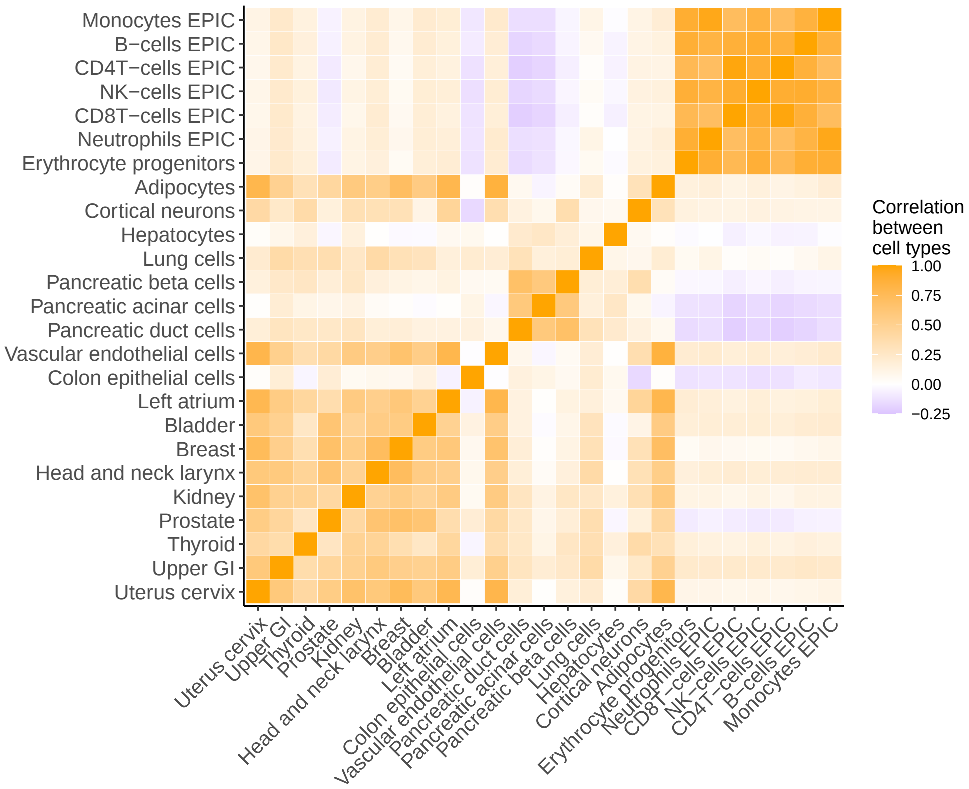


**Fig. S1 Correlation between cell types in the reference panel.** Heatmap is colored by Pearson correlation. The correlation matrix was calculated using cell type-specific CpGs only.

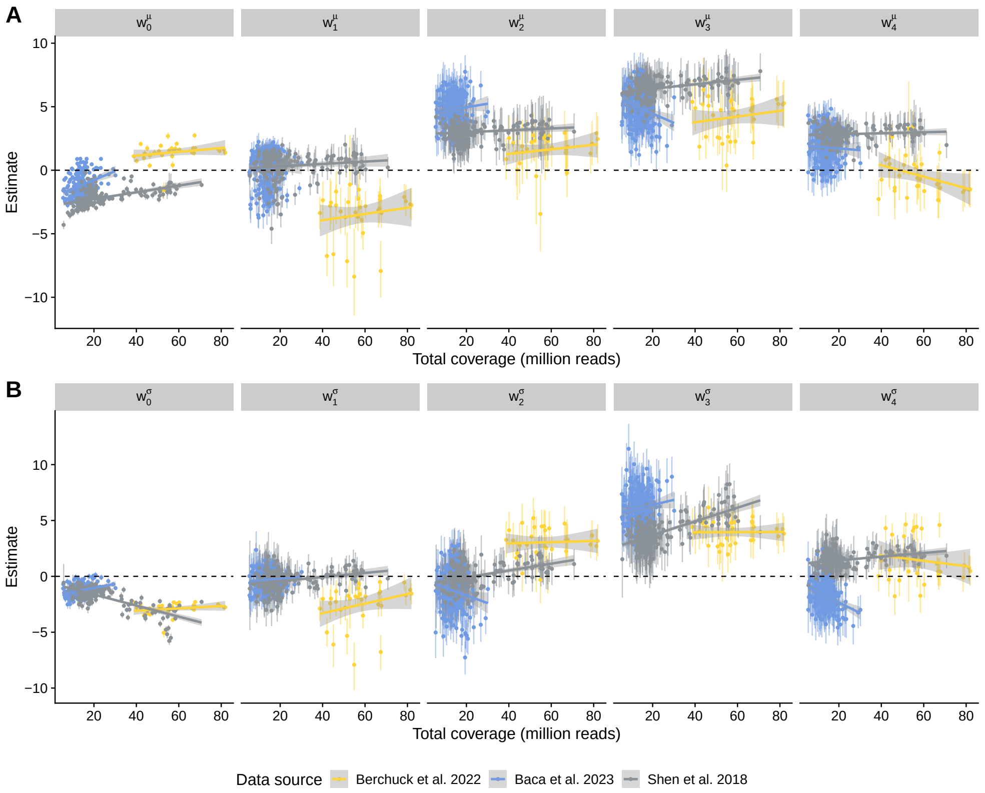


**Fig. S2 Regression coefficients in the GAM component fitted across three case studies, with respect to sample coverage levels.** Each dot represents the posterior mean of a regression coefficient, with vertical lines indicating the corresponding 95% credible intervals. Dots and intervals are colored according to data sources, representing individual samples. Solid lines and shaded areas depict the fitted linear regression models and their 95% confidence intervals for each study.

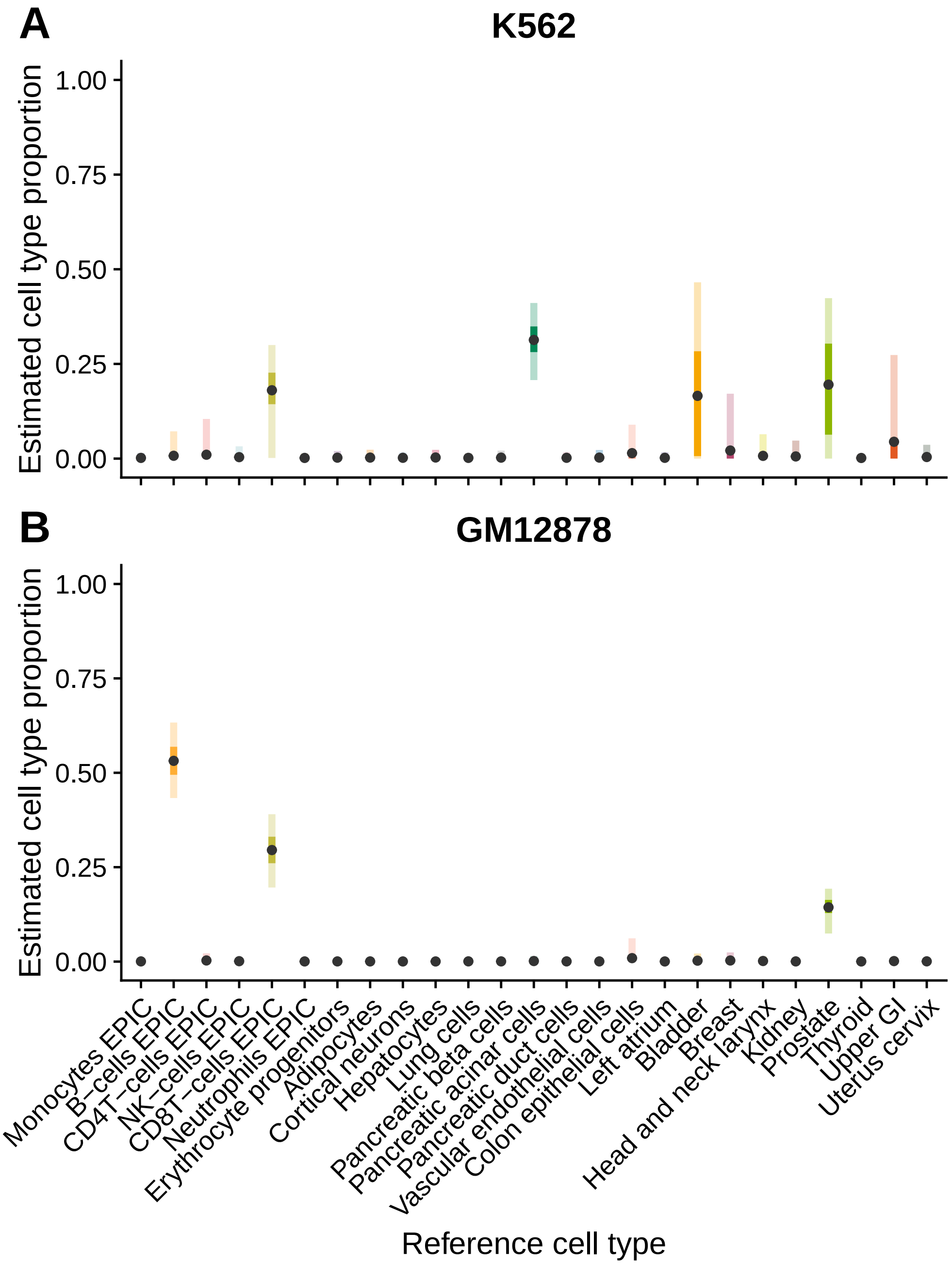


**Fig. S3 Posterior mean and credible intervals of deconvoluted proportions of all reference cell types from cell lines A) K562 and B) GM12878.** Dots indicate posterior means. The light and dark bars represent 95% and 50% credible intervals, respectively.

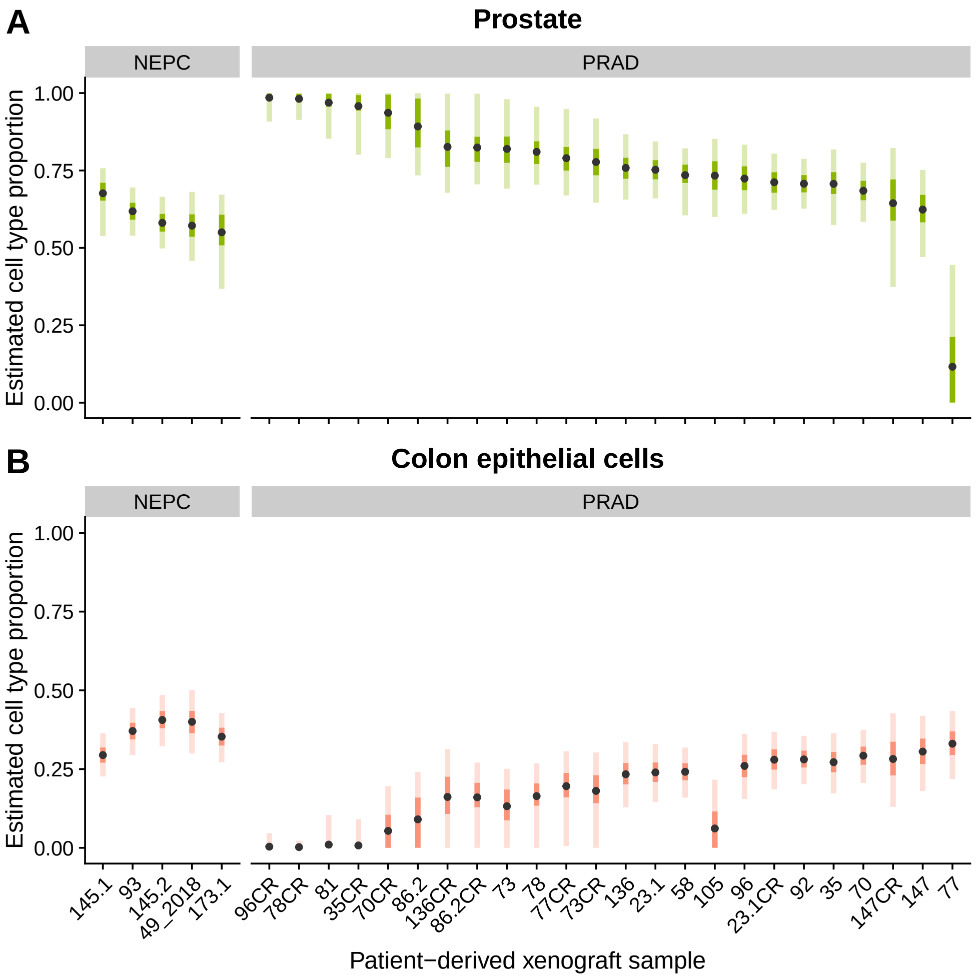


**Fig. S4 Posterior mean and credible intervals of deconvoluted proportions of prostate and colon epithelial cells in the Berchuck et al. [1] study.** Each bar represents one xenograft tissue sample. Panels correspond to two subtypes of prostate cancer, neuroendocrine prostate cancer (NEPC, n = 5) and castration-resistant prostate adenocarcinoma (PRAD, n = 24). Dots indicate posterior means. The light and dark bars represent 95% and 50% credible intervals, respectively.

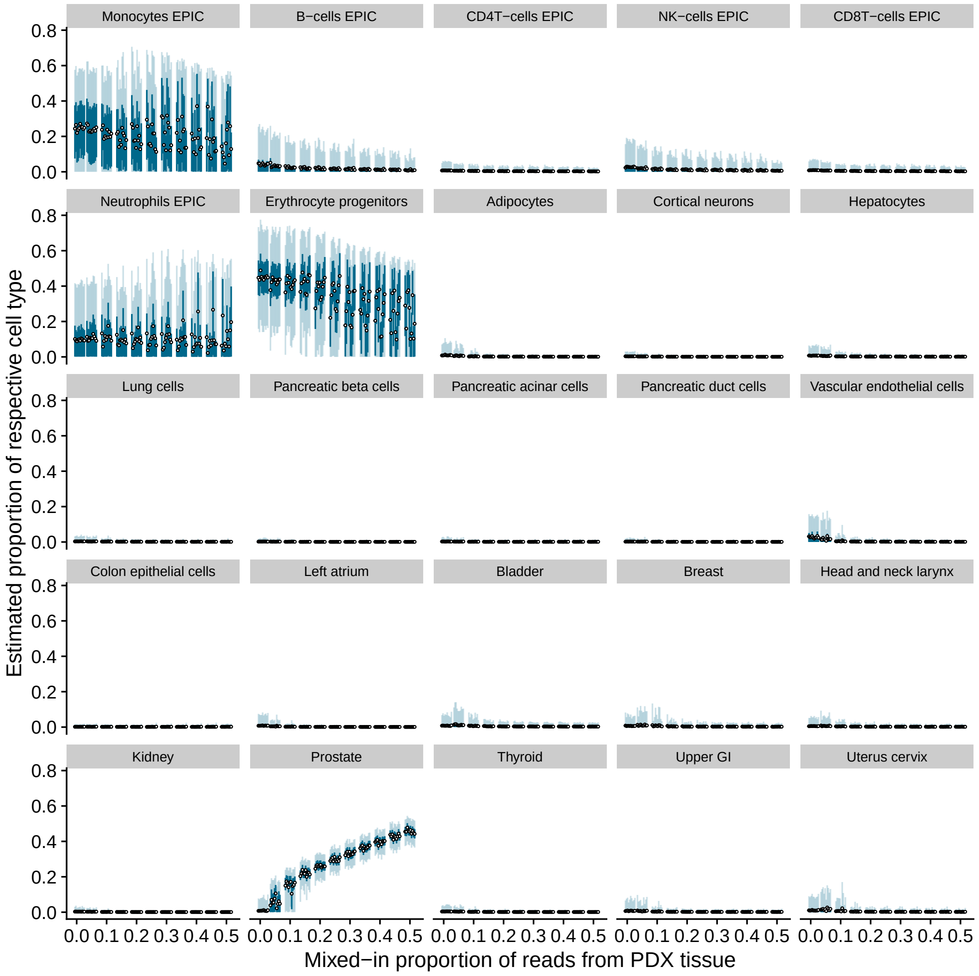


**Fig. S5 Estimated proportions of all reference cell types in PDX mixture sample pair 1.** Each panel presents results focused on a specific target cell type. Dots indicate posterior means and light and dark blue bars represent 95% and 50% credible intervals, respectively. Each posterior was fitted on a synthetic sample generated by an independent random seed.

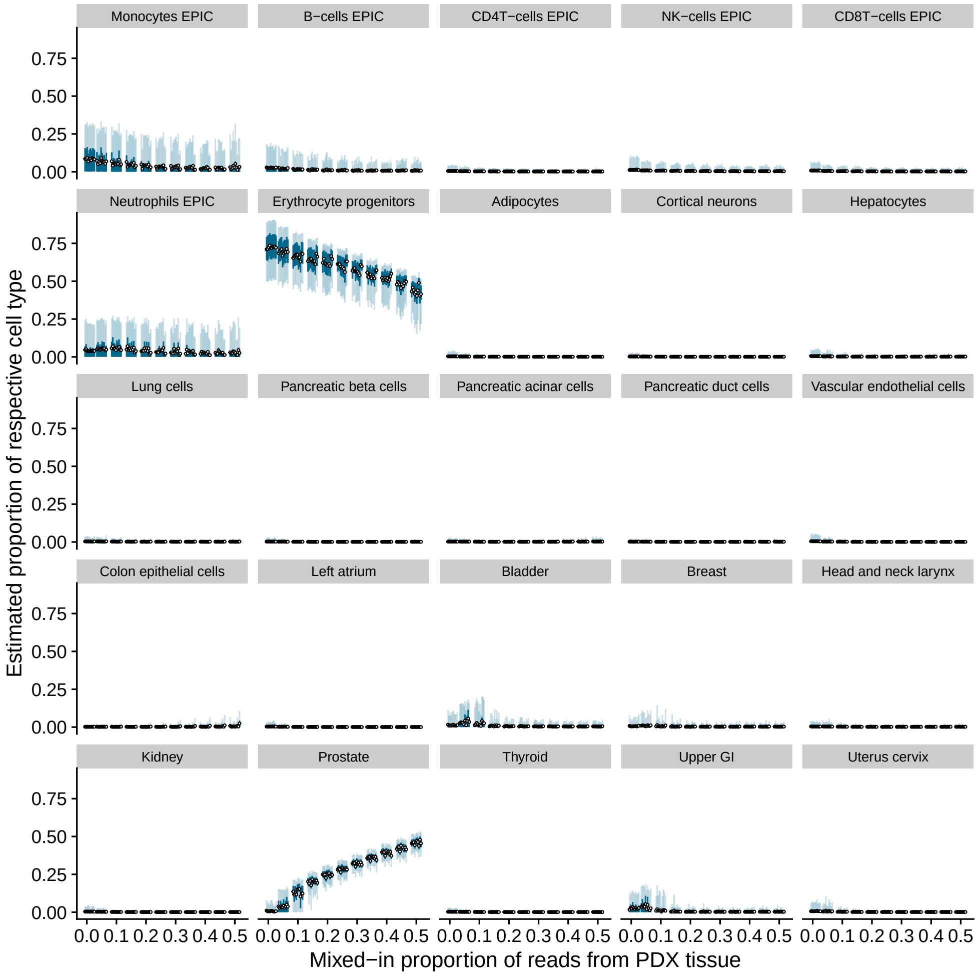


**Fig. S6 Estimated cell type proportions in PDX mixture sample pair 2.** Each panel presents results focused on a specific target cell type. Dots indicate posterior means and light and dark blue bars represent 95% and 50% credible intervals, respectively. Each posterior was fitted on a synthetic sample generated by an independent random seed.

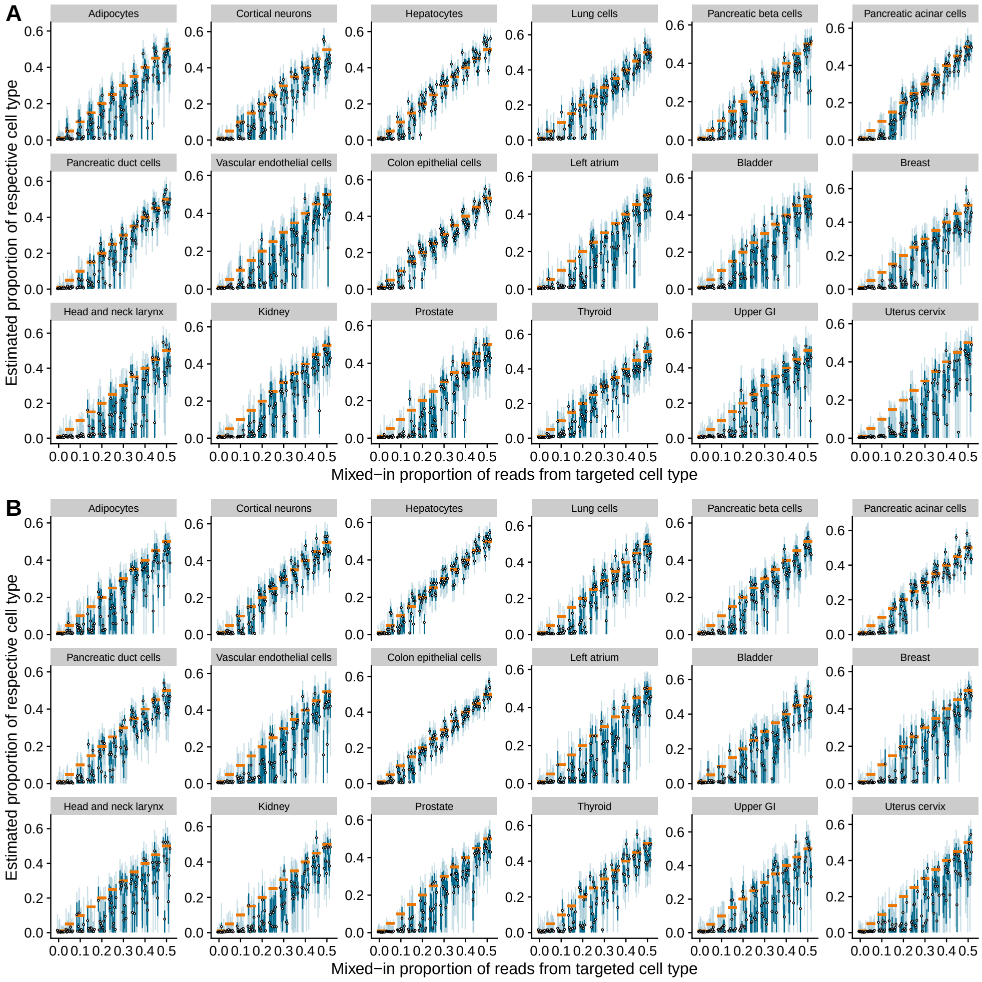


**Fig. S7 Simulation on fully synthetic data under A) low- and B) high-library size settings.** Each panel presents results focused on a specific target cell type. Orange horizontal bars represent the actual proportions of the target cell types, dots indicate posterior means, and light and dark blue bars represent 95% and 50% credible intervals respectively. Each posterior was fitted on a synthetic sample generated by an independent random seed.

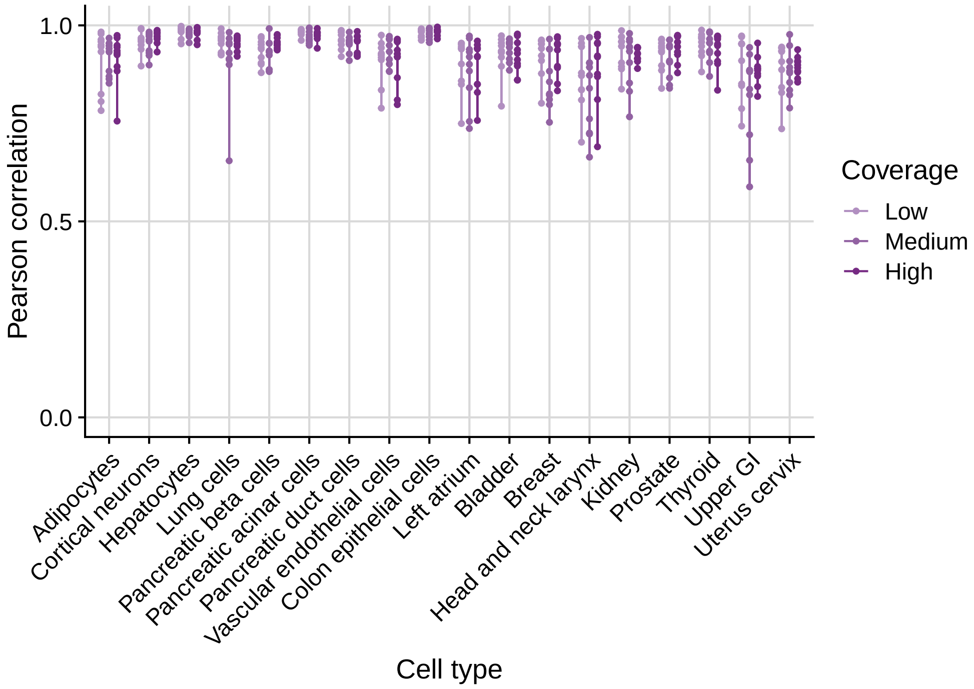


**Fig. S8 Pearson correlation between actual values and estimated values in simulation with fully model-generated data.** Each dot represents the correlation calculated from experiments with the same random seed.

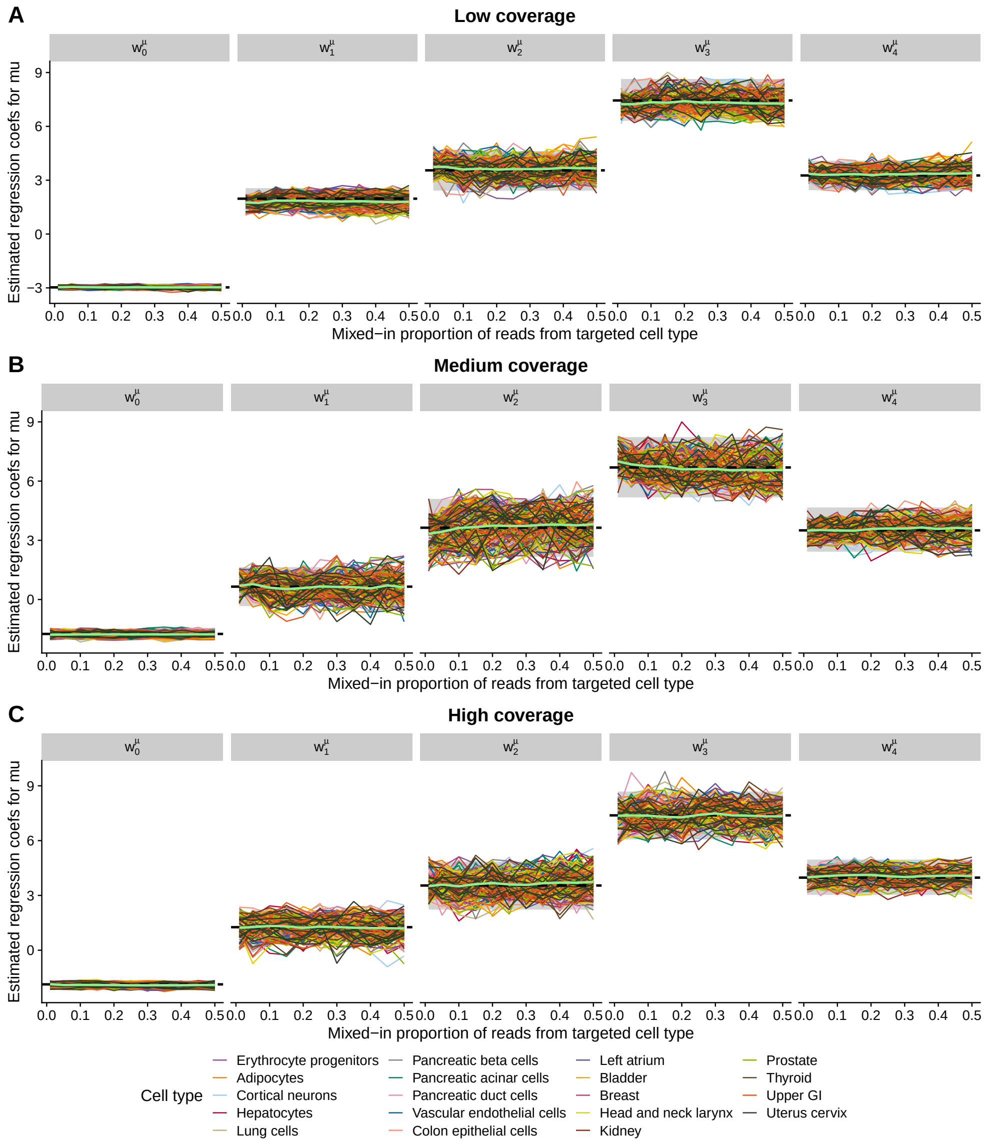


**Fig. S9 Fitted regression coefficients in $\mu(\cdot,\cdot)$ in the simulation study with fully model-generated data.** Each panel corresponds to a specific parameter. Each wiggly line illustrates the change in the estimated parameter as the mixed-in proportion of reads from a target cell type increases, with colors indicating different cell types. Bright green lines represent the average estimates across cell types, while black dashed lines denote the true parameter values. The gray shaded areas in the background indicate the 95% credible intervals for the true parameter values.

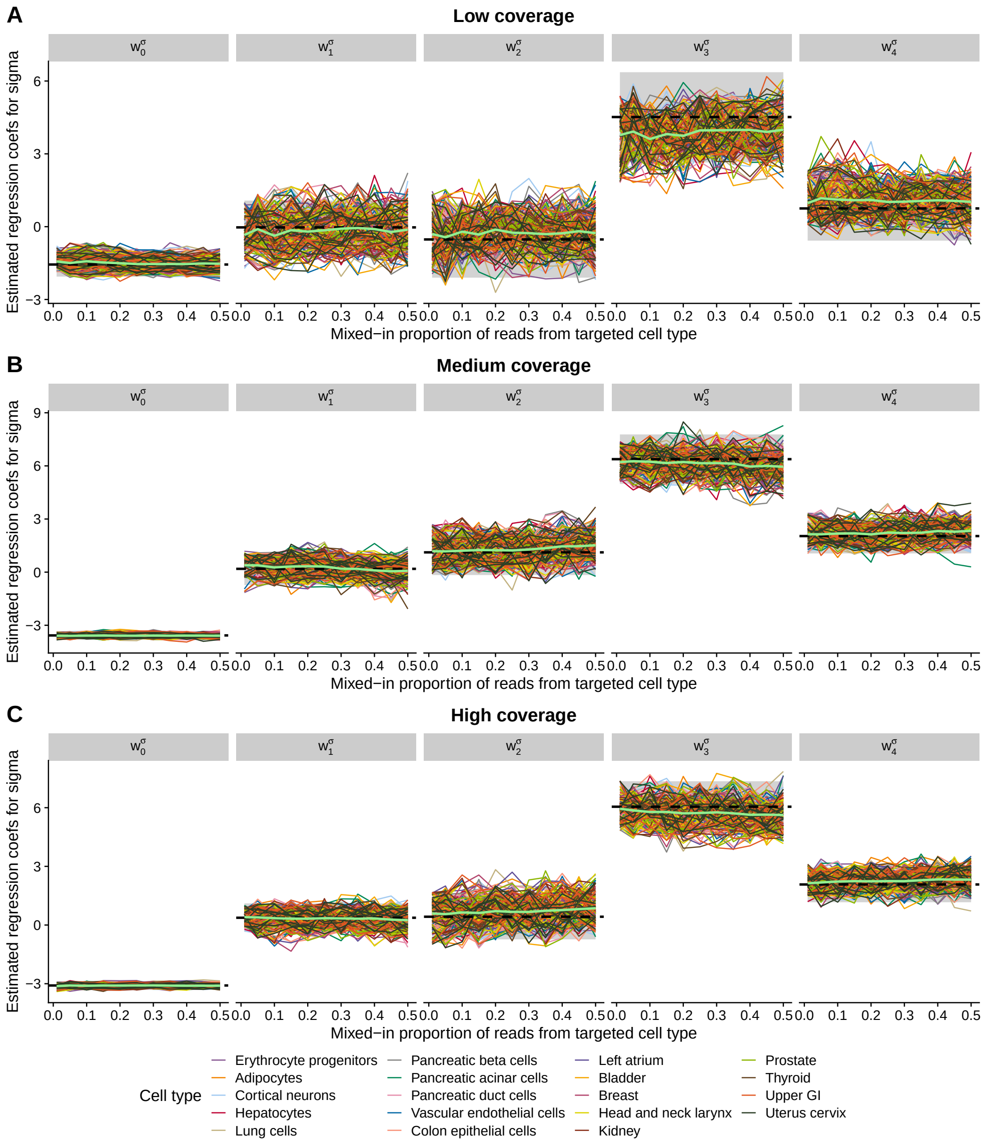


**Fig. S10 Fitted regression coefficients in $\sigma(\cdot,\cdot)$ in the simulation study with fully model-generated data.** Each panel corresponds to a specific parameter. Each wiggly line illustrates the change in the estimated parameter as the mixed-in proportion of reads from a target cell type increases, with colors indicating different cell types. Bright green lines represent the average estimates across cell types, while black dashed lines denote the true parameter values. The gray shaded areas in the background indicate the 95% credible intervals for the true parameter values.

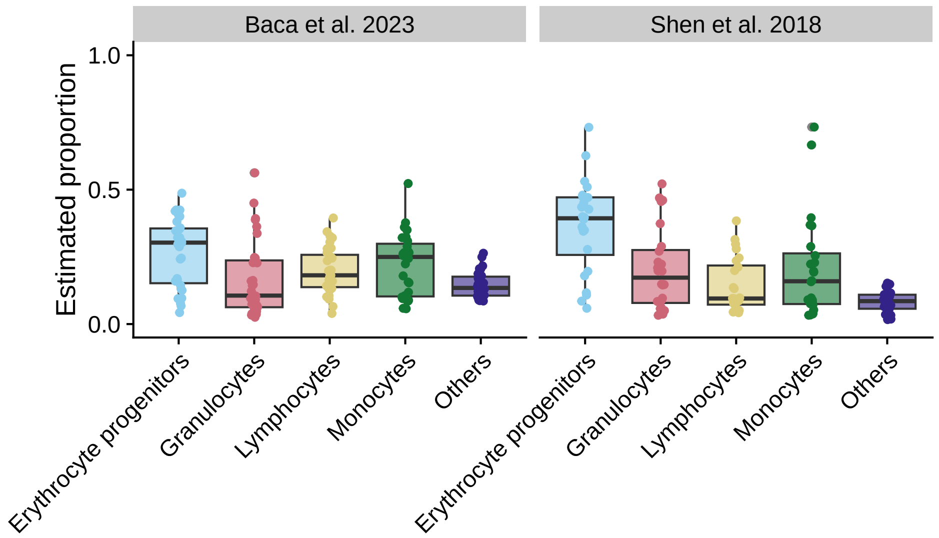


**Fig. S11 Coarse blood cell type proportions deconvoluted from healthy controls in Baca et al. [2] and Shen et al. [3].** Each dot represents an individual sample.

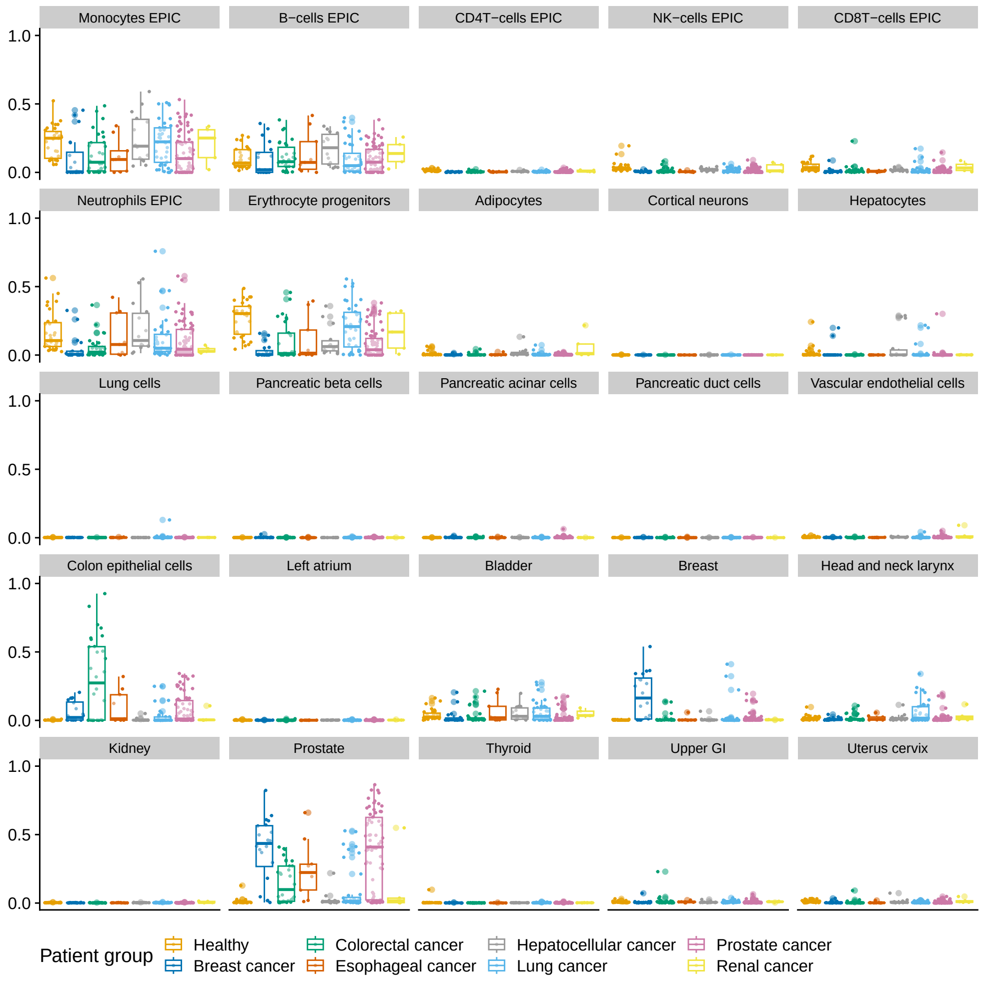


**Fig. S12 Deconvoluted proportions of all reference cell types in Baca et al. [2].** Each panel presents results focused on a specific cell type. Each dot represents an individual sample. Color represents patient groups.

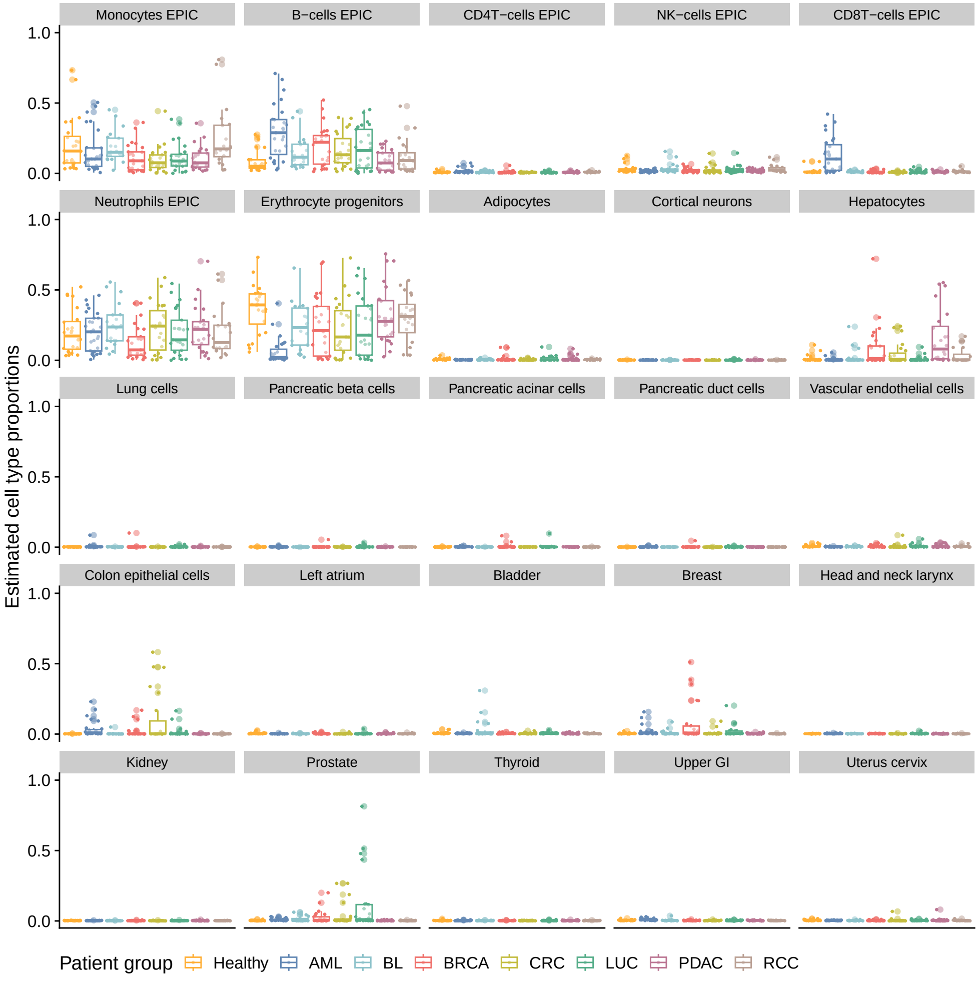


**Fig. S13 Deconvoluted proportions of all reference cell types in Baca et al. [2].** Each panel presents results focused on a specific cell type. Each dot represents an individual sample. Color represents patient groups.

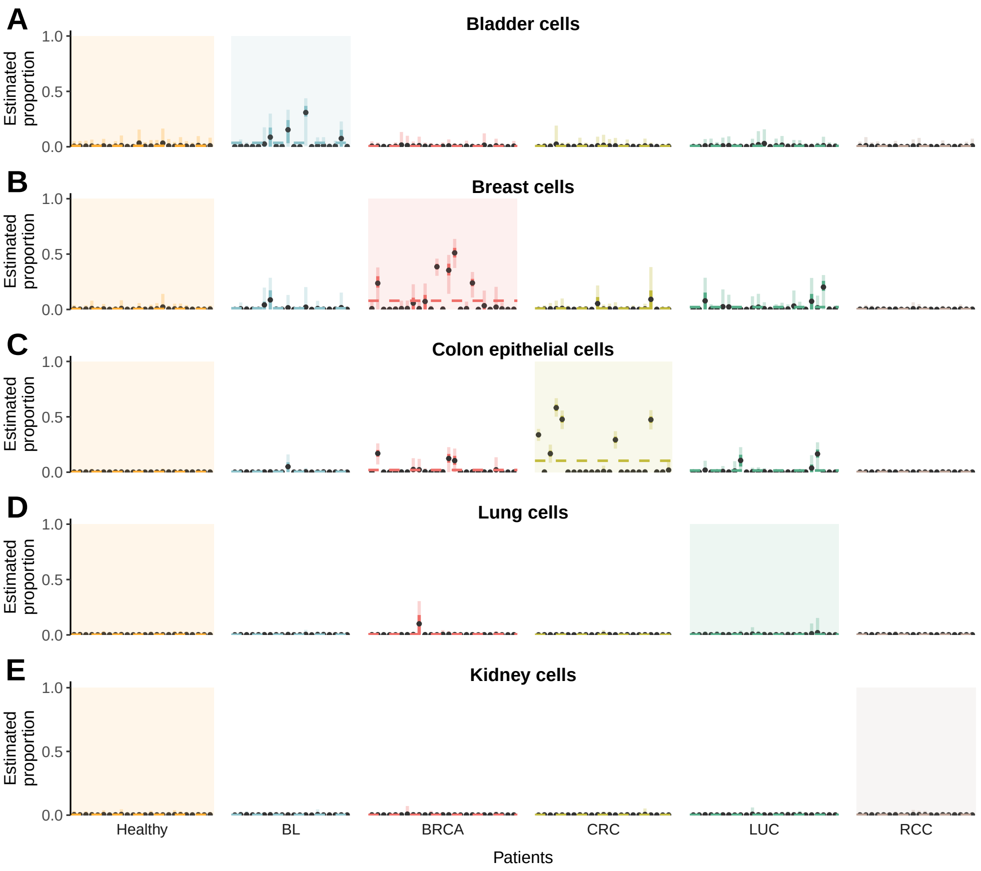


**Fig. S14 Deconvoluted cell type proportions in cancer patients in Shen et al.** [3]**.** The posterior distributions of cell type proportions for **A** bladder cells, **B** breast cells, **C**colon epithelial cells, **D** lung cells, and **E** kidney cells are presented for healthy controls and cancer patients. These five cancer types were selected because each is uniquely associated with a single cell type in the reference panel. Each interval represents the fitted posterior distribution for the cell type proportion of an individual patient, categorized and colored by patient groups. Within each interval, the dots denote the posterior mean, while the light and dark bars indicate the 95% and 50% credible intervals, respectively. Highlighted background colors emphasize the group comparisons of interest for the respective cell types. Horizontal dashed lines indicate the average posterior means of the cell type proportions within each patient group.

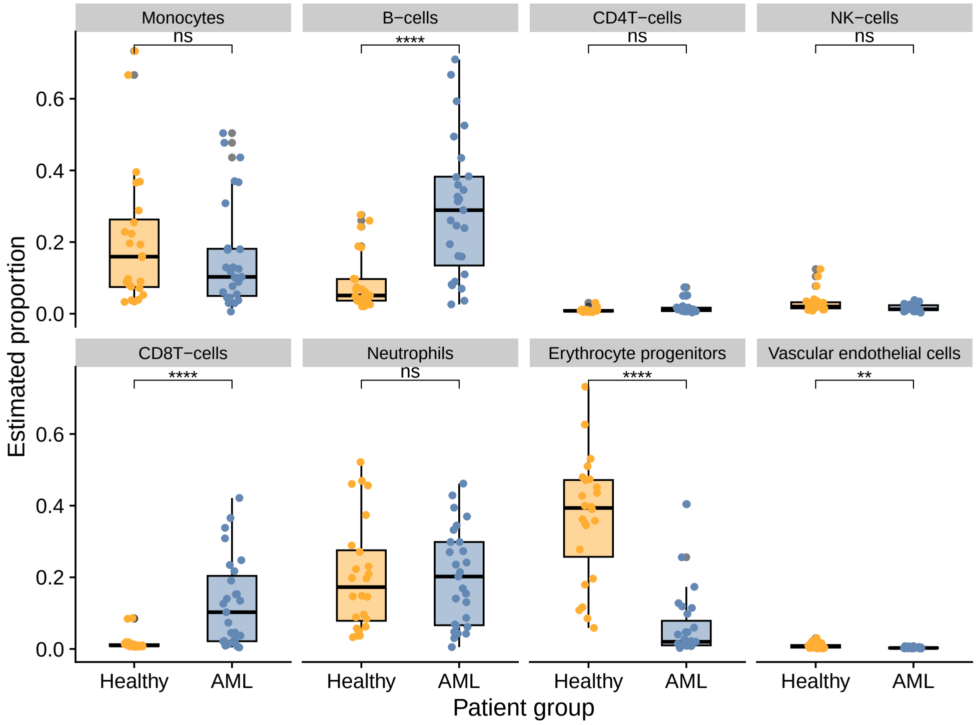


**Fig. S15 Comparison of deconvoluted blood cell types between healthy donors and acute myeloid leukemia (AML) patients from the Shen et al. [3] study.** Each panel presents results focused on a specific cell type. The number of stars represents the significance level of Bonferroni-corrected p-values from two-sample comparisons conducted via Wilcox tests ($**** :< 10^{-4}$, $*** : < 0.001$, $** :< 0.01$, $* :< 0.05$, ns: not significant).

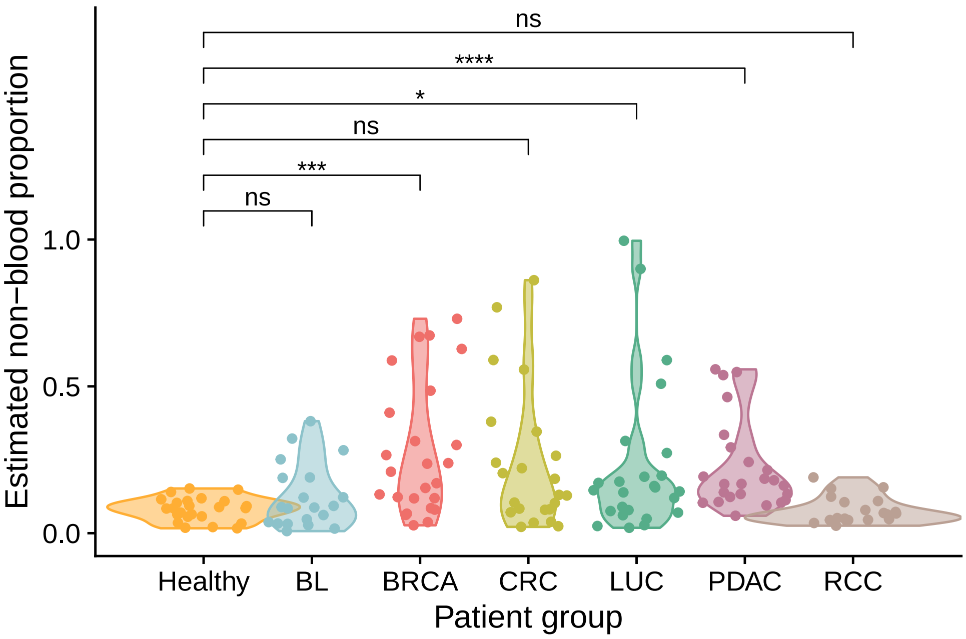


**Fig. S16 Estimated total non-blood cell proportions in all patient groups and comparison between cancer groups and the healthy controls.** The number of stars represents the significance level of Bonferroni-corrected p-values from two-sample comparisons conducted via Wilcox tests ($****: < 10^{-4}$, $***: < 0.001$, $**: < 0.01$, $*: < 0.05$, ns: not significant).

### Acronyms

*AML* acute myeloid leukemia. 16
